# The evolutionary dynamics of genetic incompatibilities introduced by duplicated genes in *Arabidopsis thaliana*

**DOI:** 10.1101/2020.09.21.306035

**Authors:** Wen-Biao Jiao, Vipul Patel, Jonas Klasen, Fang Liu, Petra Pecinkova, Marina Ferrand, Isabelle Gy, Christine Camilleri, Sigi Effgen, Maarten Koornneef, Ales Pecinka, Olivier Loudet, Korbinian Schneeberger

## Abstract

Although gene duplications provide genetic backup and allow genomic changes under relaxed selection, they may potentially limit gene flow. When different copies of a duplicated gene are pseudo-functionalized in different genotypes, genetic incompatibilities can arise in their hybrid offspring. While such cases have been reported after manual crosses, it remains unclear whether they occur in nature and how they affect natural populations. Here we identified four duplicated-gene based incompatibilities including one previously not reported within an artificial Arabidopsis intercross population. Unexpectedly, however, for each of the genetic incompatibilities we also identified the incompatible alleles in natural populations based on the genomes of 1,135 Arabidopsis accessions published by the 1001 Genomes Project. Using the presence of incompatible allele combinations as phenotypes for GWAS, we mapped genomic regions which included additional gene copies which likely rescue the genetic incompatibility. Reconstructing the geographic origins and evolutionary trajectories of the individual alleles suggested that incompatible alleles frequently co-exist, even in geographically closed regions, and that their effects can be overcome by additional gene copies collectively shaping the evolutionary dynamics of duplicated genes during population history.

## Introduction

Genetic incompatibilities describe the decrease of fitness due to incompatible allele combinations in hybrid individuals (Maheshwari and Barbash 2011). In the hybrid offspring, genetic incompatibilities result in distorted segregation of the incompatible alleles. The evolution of genetic incompatibilities has often been explained by the Bateson-Dobzhansky-Muller (BDM) model (Bateson 1909; Dobzhansky 1937; Muller 1942), where independent mutations in interacting genes get fixed in different populations, which cause deleterious epistasis and reduced fitness in their hybrids. Over the past decades, many studies have elucidated the genetic basis of such genetic incompatibilities including reciprocal pseudo-functionalization (i.e. loss of function) of duplicated genes (Fishman and Sweigart 2018; Vaid and Laitinen 2019). Gene duplications can provide genetic backup of essential genes and the basis for evolutionary novelties by allowing for new genetic and epigenetic variations (Conant and Wolfe 2008; Kondrashov 2012; Panchy et al. 2016). However, in some cases pseudo-functionalization of duplicated essential genes may occur independently in both copies in different individuals. This in turn can lead to the loss of any functional gene copy in hybrid offspring of such individuals, and thereby cause severe genetic incompatibilities (Lynch and Force 2000).

Genetic incompatibilities introduced by duplicated genes have been reported within inter/intra-specific hybrids of *Arabidopsis thaliana* (Bikard et al. 2009; Durand et al. 2012; Agorio et al. 2017), rice (Mizuta et al. 2010; Yamagata et al. 2010; Nguyen et al. 2017) or *Mimulus* (Zuellig et al. 2017). Identification of these incompatible alleles, however, often relied on genetic mapping in experimental populations, which is a time consuming and costly process. As incompatible alleles are frequently introduced by loss-of-function (LoF) (epi)mutations (Bikard et al. 2009; Blevins et al. 2017), initial examination of LoF (epi)mutations within whole-genome sequence data could be a shortcut to quickly target promising candidates.

Although several incompatible alleles from duplicated genes have been identified in *A. thaliana*, it is still unclear how these incompatible alleles originate and evolve in natural populations, and how the populations adapt to the reduction in fitness. Untangling the complex evolutionary process would require accurate (epi)genotypes of incompatible genes across sufficiently large natural populations. The Arabidopsis 1001 Genomes Project (Alonso-Blanco et al. 2016) and 1001 Epigenomes Project (Kawakatsu et al. 2016) have released substantial omics data, which can be used to unravel the evolutionary trajectory of such incompatible alleles.

Here, we created an extended version of the Arabidopsis multiparent RIL population (Huang et al. 2011) to identify genetic incompatibilities between several different genotypes simultaneously. Based on distorted segregation of duplicated genes, we mapped four genetic incompatibilities. Unexpectedly, however, we identified several, healthy RILs which carried presumably incompatible allele combinations. Further analysis of their genomes revealed additional gene copies rescuing these severe incompatibilities. Encouraged by this, we searched for incompatible allele combinations within 1,135 accessions of the 1001 Genomes Project (Alonso-Blanco et al. 2016), where these combinations were surprisingly common. Using the incompatible allele combinations as phenotypes we mapped modifiers of all four incompatibilities using GWA. The loss-of-function alleles from duplicated genes were geographically widely distributed, and co-existed with additional gene copies in the same regions showing how additional gene copies can overcome differential copy loss in a population.

## Results

### Identification of incompatible gene pairs within an intercross population

We used the Arabidopsis Multiparent RIL (AMPRIL) population to find incompatible alleles that arose from duplicated genes. The eight AMPRIL founder accessions (An-1, C24, Col-0, Cvi-0, Eri-1, Kyo, L*er* and Sha) were selected across the entire geographic distribution of *A. thaliana* including the Northern hemisphere and the Cape Verde Islands. Recently, we generated chromosome-level genome assemblies of all seven, non-reference founder genomes (Jiao and Schneeberger 2020). The first release of the AMPRIL population (AMPRIL I) contained six subpopulations (referred to as ABBA, ACCA, ADDA, BCCB, BDDB, CDDC) derived from reciprocal diallel crosses between four hybrids (A: Col-0 x Kyo, B: Cvi-0 x Sha, C: Eri-1 x An-1, D: L*er* x C24) and the subsequent selfing to the F4 generation by single-seed descent (Huang et al. 2011). Here we present the extension of the AMPRIL population with six new subpopulations referred to as EFFE, EGGE, EHHE, FGGF, FHHF and GHHG (E: Col-0 x Cvi-0, F: Sha x Kyo, G: Ler x An-1, H: Eri-1 x C24) based on different diallel intercrossing scheme and selfing of the recombinant genomes until the F6 generation (Fig. 1a). Each subpopulation consists of approximately 90 individuals representing recombinants of four founders. In total, 992 RILs from all twelve subpopulations were sequenced and analyzed using RAD-seq (Baird et al. 2008) (Supplementary Data 1) and genotyped with ∼2 million high-quality SNP markers. We used a Hidden Markov Model to reconstruct the parental haplotypes (identity-by-descent) including residual heterozygous regions (Rowan et al. 2015) (Supplementary Fig. 1 and Supplementary Note 1). The genotyping resulted in 12,878 different recombination breakpoints (on average one breakpoint per 9.3 kb) across the entire population. This allowed us to divide the genome of each progeny into 12,883 haplotype blocks, where each block relates to the haplotype(s) of only one (homozygous regions) or two (heterozygous regions) of the founder haplotypes.

**Figure 1.**
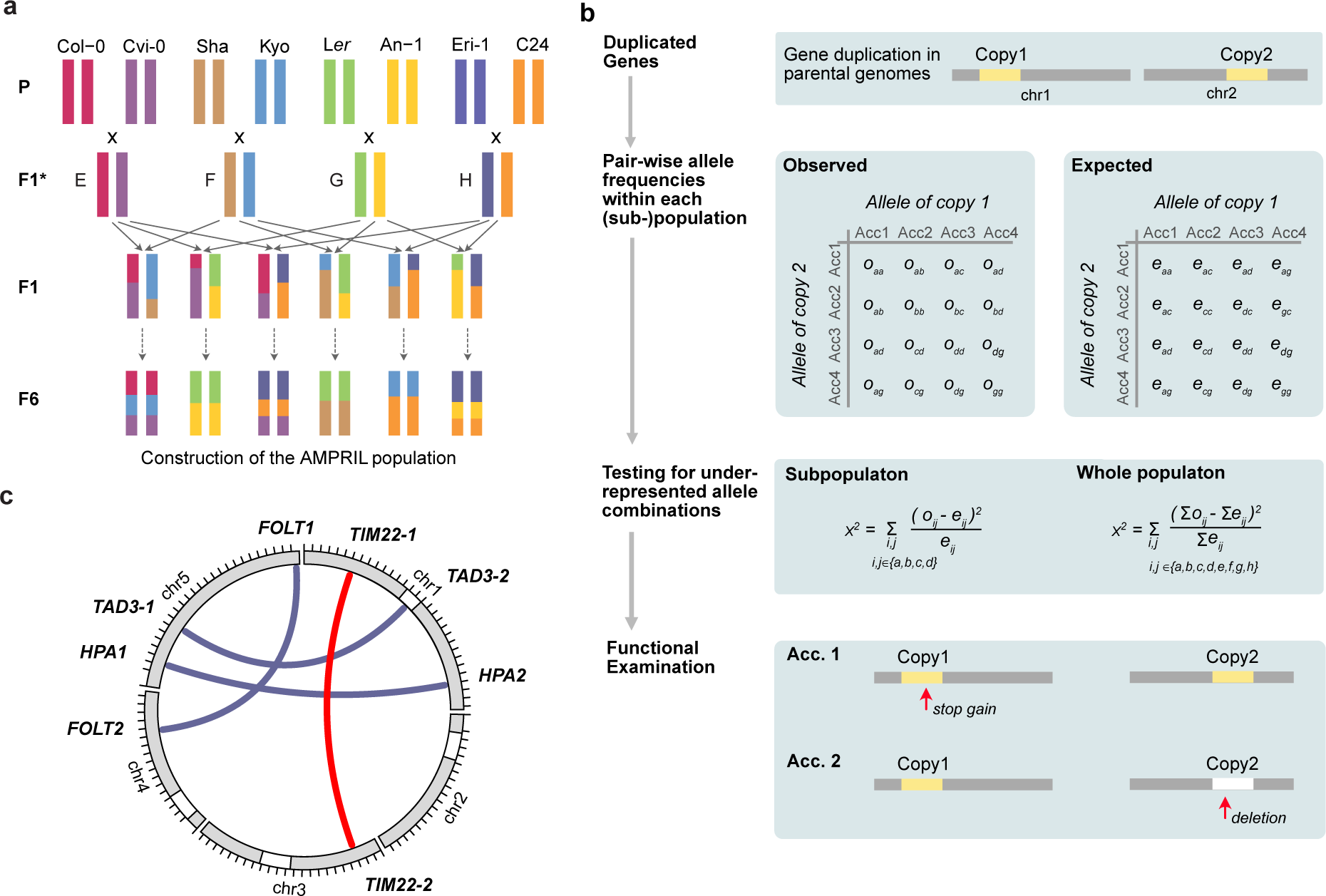
Identification of genetic incompatibilities introduced by duplicated genes in an intercross population. (**a**) Construction of the extended AMPRIL population. Eight different *A. thaliana* accessions were used as founder lines. For each of the six subpopulations, two F1* hybrids, which were generated by crossing two founder lines, were again crossed to give rise to the F1 individuals of each population. The F1 individuals were further self-crossed to the F6 generation. (**b**) Workflow for the identification of potentially incompatible alleles in duplicated genes. Unlinked (i.e. on separate chromosomes) duplicated genes were selected and the expected and observed frequencies of all haplotype combinations between the two copies were calculated for each subpopulation and the whole population. Under-represented allele combinations were identified using χ^2^ test. Each gene duplication with significantly underrepresented allele combinations was evaluated for non-functionalized or deleted gene copies in the respective parental genomes. (**c**) The location of incompatible alleles in four duplicated gene pairs identified in the AMPRIL population including one so-far unknown incompatibility (red) and one detected a posteriori in an informed way (*TAD3*).

We developed a two-step workflow to combine genetic and genomic evidence to quickly identify incompatible alleles of duplicated genes (Fig. 1b). In the first step, we selected 781 distal (inter-chromosome) duplicated gene pairs including 612 gene pairs in which the reference sequence contains two copies and other founder genomes feature at least one copy (Supplementary Data 2). In the remaining 169 gene pairs, the reference sequence only has one copy and at least one other parental genome has an additional copy in a different chromosome. As genetic incompatibility leads to the underrepresentation of incompatible allele combinations (Ackermann and Beyer 2012; Corbett-Detig et al. 2013), we searched for significant distortions from the expected frequencies of all parental allele combinations across all 781 duplicated gene pairs in all twelve subpopulations, two merged subpopulations (ABBA and EFFE, CDDC and GHHG as they share the same founders), and the whole population (see Methods). These tests revealed significant distortions in 236 gene pairs (χ^2^ test, p-value < 0.05, multiple testing corrected) in at least one of the populations.

However, the observed distortions do not necessarily result from genetic incompatibilities in the tested gene. Alternatively, such distortions can also occur if the tested gene duplicate is closely linked to a genetic incompatibility. Hence, in a second step, we examined the alleles of the gene pairs in the founder genomes for loss-of-function (LoF) variations or hypermethylated promoters (Fig. 1b, see Methods). This examination revealed three gene pairs with functional disruption in both of the duplicates in at least one of the founder genomes. These three duplicated genes included two, *HISTIDINOL PHOSPHATE AMINOTRANSFERASE* (*HPA*) (Bikard et al. 2009) and *FOLATE TRANSPORTER* (*FOLT*) (Durand et al. 2012), which were already known for their ability to introduce genetic incompatibilities, as well as one gene pair, which so-far was not reported as the genetic basis for a genetic incompatibility, *TIM22* (*TIM22-1*: *AT1G18320, TIM22-2*: *AT3G10110*) (Fig. 1c). For all other 233 gene pairs we could not identify non-functional alleles in both copies.

We noted that another duplicated gene, *tRNA ADENOSINE DEAMINASE 3* (*TAD3)*, which is also know to introduce a genetic incompatibility (Agorio et al. 2017), was not considered by our initial testing even though the genotypes of the AMPRIL founders should lead to incompatible allele combinations in the RIL populations: all founder genomes except for Kyo have a functional *TAD3-1* (Supplementary Table 1), while the Kyo *TAD3-1* gene is silenced most likely due to its methylated promoter (similar to the Nok-1 and Est-1 accessions in which the incompatibility was described originally (Agorio et al. 2017)). The lack of a functional *TAD3-1* in Kyo is counterbalanced by additional copies (*TAD3-2*) (which however were not part of the main chromosome scaffolds of the Kyo genome assembly). Therefore, the *TAD3* gene duplicate was not considered initially, however, we also used this incompatibility for further analysis.

### Genetic incompatibility introduced by diverged copies of *TIM22*

Before we started our analysis of the four incompatibilities, we verified that the LoF alleles of *TIM22* are in fact the causal basis of the genetic incompatibility that we observed in the AMPRIL population. *TIM22* encodes for a mitochondrial import inner membrane translocase subunit of the *TIM17*/*TIM22*/*TIM23* family protein (Murcha et al. 2007). All eight founders feature two annotated *TIM22* copies, while Cvi-0 includes an extra truncated copy ∼31 kb downstream to *TIM22-2* (Fig. 2a, b). We found significant segregation distortions in the complete AMPRIL population and one subpopulation, EGGE, where the double homozygous allele combination *TIM22-1*^Col-0^*TIM22-2*^Cvi-0^ was significantly underrepresented (Supplementary Table 2 and 3).

**Figure 2.**
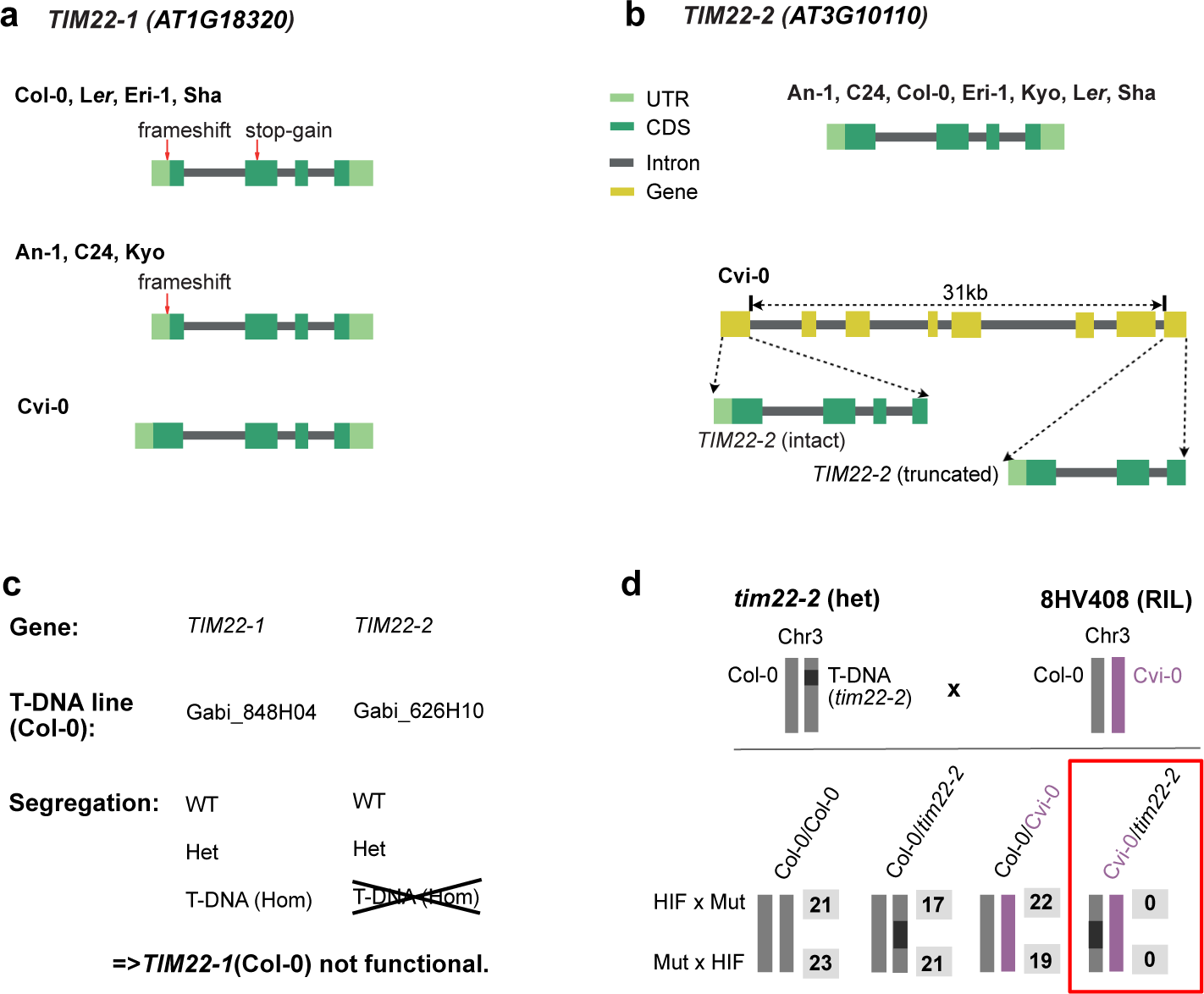
Genomic and genetic evidence of incompatible *TIM22* alleles. **(a)** Gene structure of *TIM22-1* in the genomes of the eight AMPRIL parents. The loss-of-function variants (1bp frameshift indel and a premature stop-codon) relative to the intact *TIM22-1*^Cvi-0^ are shown. The deleterious effect of the frameshift is erased by an alternative translation start site. **(b)** Gene structure of *TIM22-2* in the genomes of the eight AMPRIL parents. The eight accessions share the structure of *TIM22-2* without recognizable loss-of-function variants, however, a truncated copy of *TIM22* could be found in Cvi-0 ∼31 kb downstream of *TIM22-2*. **(c)** Segregation of T-DNA alleles within the descendance of 2 segregating Col-0 T-DNA mutant lines (*tim22-1* and *tim22-2*). **(d)** A heterozygous T-DNA line (*tim22-2*) in the Col-0 background (i.e. non-functional for *TIM22-1*) was crossed to 8HV408 (heterozygous Col-0/Cvi-0 at *TIM22-2* and homozygous for the Col-0 allele at *TIM22-1* i.e. nonfunctional for *TIM22-1*). The number of all four possible F1 progenies are shown (in grey) for both cross directions. While the *TIM22-2*^*Col-0*^ allele can complement the T-DNA, the *TIM22-2*^*Cvi-0*^ could not, implying that the *TIM22-2*^*Cvi-0*^ allele is non-functional. HIF: heterogeneous inbred families, Mut: mutant.

We observed an in-frame premature stop-codon in Col-0 (mis-annotated in the reference annotation) suggesting that *TIM22-1* is not functional in Col-0 (Fig. 2a and Supplementary Table 4). To test if *TIM22-1* is truly non-functional in Col-0, we used the segregation of two T-DNA insertion mutants in the two *TIM22* paralogs in Col-0. This showed that *tim22-1* could be homozygous for the T-DNA insertion allele but the T-DNA in *tim22-2* could not be found in homozygous state (Fig. 2c). This suggests that, in Col-0, *TIM22-1* is not functional and *TIM22-2* is the only functional copy.

The *TIM22* paralogs co-located within the regions of a previously reported genetic incompatibility (hereafter named as LD2: LD2.1 for the locus at chromosome 1 and LD2.3 for the locus at chromosome 3) which was mapped in a Cvi-0 x Col-0 RIL population (Simon et al. 2008). The genetic underpinnings of this incompatibility however were still unknown. This incompatibility was expressed by a striking underrepresentation of homozygous LD2.1^Col-0^ combined with homozygous LD2.3^Cvi-0^, which was in agreement with the reduced allele combinations in the AMPRIL population (Supplementary Table 2 and 3). Therefore, we generated heterogeneous inbred family (HIF) lines from Cvi-0 x Col-0 RILs to fine-map LD2.1 and LD2.3 to respectively 70kb and 34kb intervals (Supplementary Fig 2). The two candidates *TIM22-1* and *TIM22-2* remained within the intervals.

To validate their causative role, we conducted a complementation cross between a heterozygous T-DNA mutant in *TIM22-2* in a Col-0 background (i.e. *TIM22-1* was non-functional) and the original HIF line in which *TIM22-1* was homozygous for the Col-0 genotype (i.e. also non-functional) and *TIM22-2* was heterozygous for Col-0/Cvi-0 (Fig. 2d). Within 123 hybrids of the offspring, among the four possible allelic combinations at LD2.3, we did not find any hybrids combining a Cvi-0 and a T-DNA alleles at *TIM22-2* (Fig. 2d), providing strong genetic evidence that the Cvi-0 allele cannot complement a knockout (T-DNA) allele at *TIM22-2* (in a background without other functional *TIM22* allele) and is thus non-functional.

Collectively, these segregation and complementation crosses show that the combination of different non-functional alleles of the *TIM22* copies leads to a drastic reduction of allelic combinations in offspring populations, and thus evidence the causative role of *TIM22* in this genetic incompatibility.

### Natural modifiers can rescue incompatible allele combinations

Incompatible allele combinations can result in severe phenotypic defects, which lead to the reduction or the full absence of specific allele combinations. For example, the double homozygous non-functional allele combination of *HPA1/HPA2* results in embryo lethality (Bikard et al. 2009) and thereby wipes out all carriers of the incompatible allele combination. *HPA* encodes a histidinol-phosphate amino-transferase for the biosynthesis of histidine, an essential amino acid (Muralla et al. 2007). All eight AMPRIL founders except of Cvi-0 have a functional *HPA1* and a non-functional *HPA2* due to a premature stop codon or a hypermethylated promoter, while Cvi-0 carries a functional *HPA2* allele, but does not carry *HPA1* at all (Supplementary Table 5). Unexpectedly, however, we did observe homozygous *HPA1/HPA2* incompatible allele combinations (*HPA2*^*−/−*^*HPA1*^*−/−*^) in eleven of the AMPRIL lines within the ABBA and EFFE subpopulations which were derived from Col-0, Cvi-0, Kyo and Sha (Supplementary Table 6-8). Further analysis of these populations revealed an extremely high frequency of the Kyo allele on chromosome 4 (Supplementary Fig. 3), making us recognize that the eleven AMPRIL lines with the incompatible allele combinations all carried at least one Kyo allele at chr4:9.2-13.7 Mb (Fig. 3a). This suggested that Kyo might contain a modifying allele of the incompatibility complementing the lethal allele combination in this region (a similar feature was described as “conditional incompatibility” by Bikard et al. when analyzing crosses between Jea and Col-0 (Bikard et al. 2009)). Indeed, when we checked the long-read assembly of the Kyo genome, we identified an additional *HPA* copy (hereafter named as *HPA3*) which co-located with the mapping interval at chr4:11.11 Mb (Fig. 3b), while we could not find this allele in any of the other AMPRIL founder genomes.

**Figure 3.**
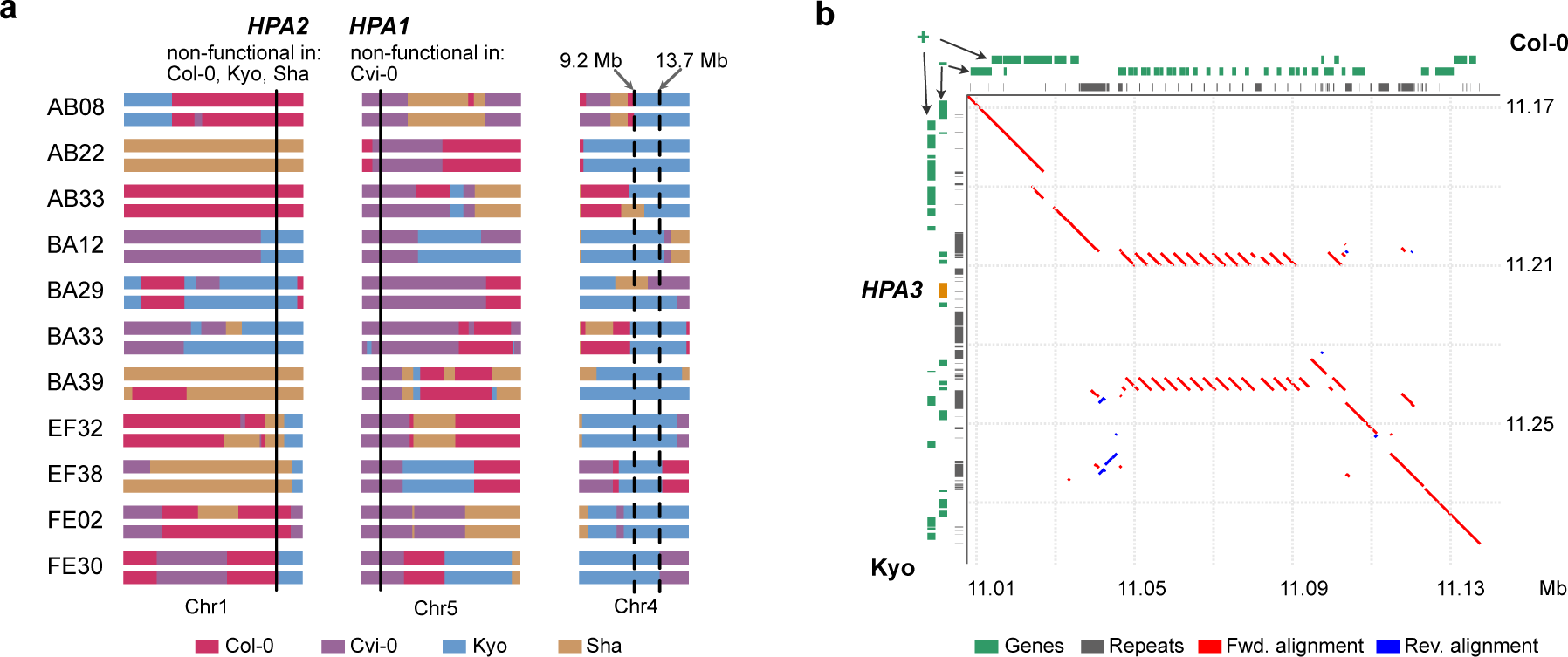
Incompatible allele combinations of *HPA* rescued by an additional gene copy. **(a)** Schematic of the genomes of 11 AMPRILs with presumably incompatible allele combinations of *HPA1* and *HPA2*. All these genotypes carry at least one Kyo allele at chr4: 9.2 – 13.7 Mb, suggesting that a Kyo allele in this region can rescue the incompatibility. **(b)** Sequence alignment around the *HPA3* locus on chromosome 4 between Col-0 and Kyo. The position of the third *HPA3* copy in Kyo is marked in orange. Red line: forward alignment; blue line: reverse alignment. Genes arrangement at forward (+) and reverse (-) strands, and repeat annotations are shown at the top (Col-0) or left (Kyo) axes.

### Mapping modifiers of incompatible allele combinations in natural populations

This analysis showed that the negative effects of incompatible allele combinations can be overcome if they are rescued by modifying alleles. Therefore, the virtual absence of functional alleles (among the know loci) of a duplicated gene could act as a molecular phenotype to map the location of additional (rescuing) alleles by genome-wide association (GWA) mapping even in natural populations.

To do this, we searched for incompatible allele combinations (separately for all four incompatibilities) in each of the 1,135 accessions of the 1001 Genomes Project (Alonso-Blanco et al. 2016) and used these allele combinations as phenotype for a GWA (Fig. 4). To define non-functional alleles, we used the resequencing data to search for LoF variations, and the methylome data from the 1001 Epigenomes Project (Kawakatsu et al. 2016) to identify methylated (silenced) promoters (Fig. 4a, see Methods). Additionally, RNA-seq data were explored to distinguish pseudo-heterozygous variants (which exist due to inaccurate short-read alignment at duplicated genes) and to check for gene silencing.

**Figure 4:**
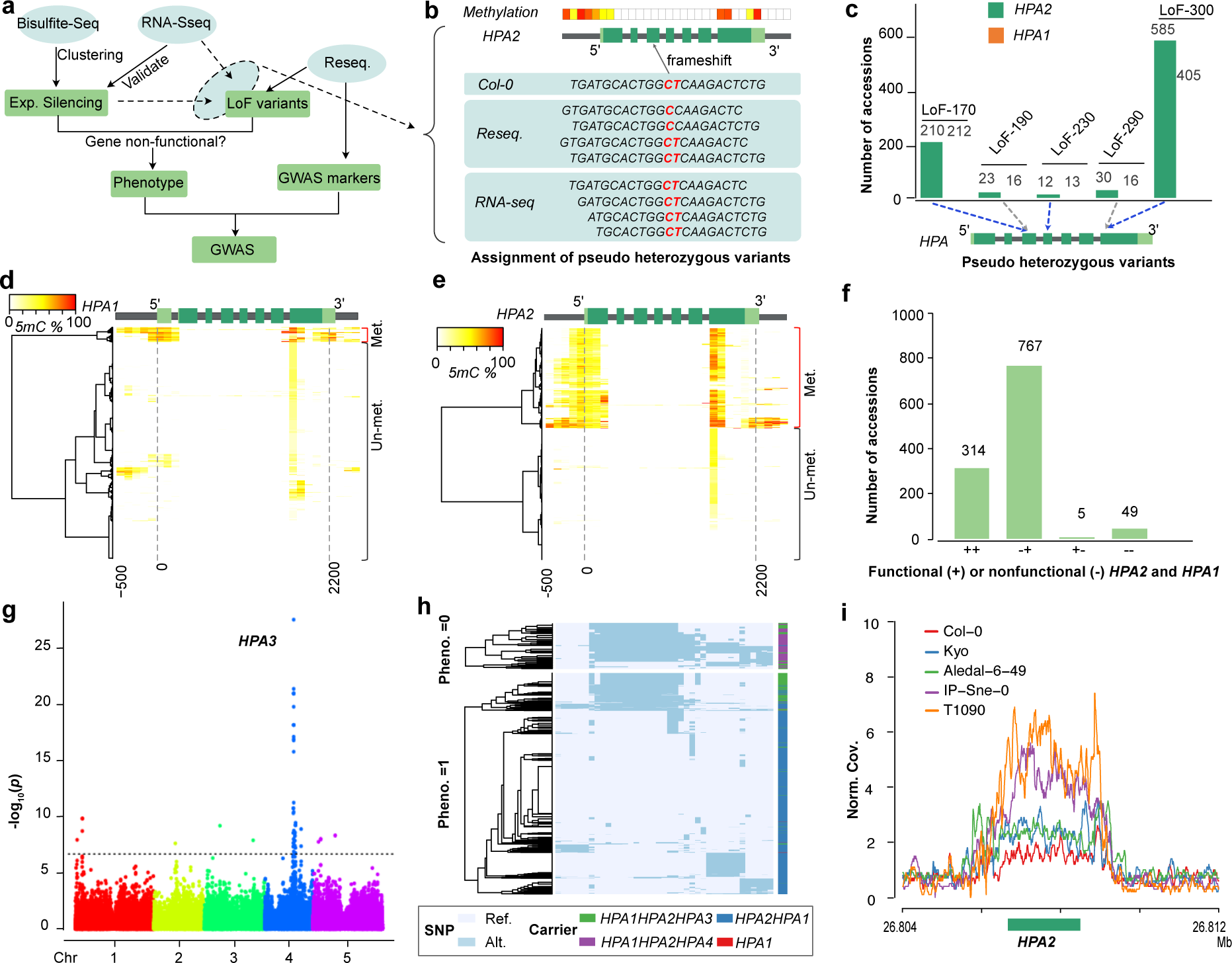
Mapping natural modifiers of genetic incompatibilities using GWAS. **(a)** Workflow for the identification of modifying alleles of genetic incompatibility using GWAS. **(b)** Schematic example to illustrate how genome-wide methylation and RNA-seq data are used to assign pseudo-heterozygous variants to a specific gene copy despite the repetitive nature of the short-read alignments within duplicated genes. If a specific variation is present in DNA data, but absent in RNA data and one of the gene copies is methylated (i.e. likely expression silenced) the pseudo-heterozygous variation is assigned to this (expression silenced) gene copy. Light green: untranslated region, green: coding region, gray: intron or gene up/down-stream. **(c)** Pseudo-heterozygous LoF variants found in the short read alignments (of 1,135 *A. thaliana* genomes(Alonso-Blanco et al. 2016)) at *HPA2* and *HPA1*. LoF-170, LoF-230, and LoF-300 could be assigned to *HPA2* (using the procedure of (b)), while LoF-190 and LoF290 could not be assigned to either of the gene copies. **(d, e)** Hierarchical clustering of DNA methylation profiles in *HPA1* (d) and *HPA2* (e) based on the methylomes of 888 *A. thaliana* accessions from the 1001 Epigenome Project (Kawakatsu et al. 2016) (NCBI GEO accession: GSE43857). Methylation profiles calculated within 100 bp sliding windows from 500 bp upstream of the transcription start site to 300 bp downstream of the transcription end site. **(f)** The number of accessions with different functional copies of *HPA*2 and *HPA1* across 1,135 *A. thaliana* accessions. **(g)** Manhattan plot of a GWA using the absence of any functional *HPA* gene as phenotype. The most significantly associated locus reveals the region of *HPA3*. The dashed line indicates the significant threshold after multiple testing correction (−log_10_(*P*)=6.68). **(h)** Two heatmaps of haplotype clustering (defined by the 39 significantly associated markers around the *HPA3* locus) shown for accessions with (below) or without (up) functional copies of *HPA2* or *HPA1*. **(i)** The normalized short read mapping coverage (Norm. Cov.: average mapping coverage at *HPA2* divided by average mapping coverage at whole genome) around *HPA2* based on read mappings against the Cvi-0 genome including only one *HPA* gene. (Data from five accessions are shown as examples to illustrate patterns of different *HPA* copies: Col-0: *HPA2HPA1*; Kyo and Aledal-6-49: *HPA2HPA1HPA3*; IP-Sne-0 and T1090: *HPA2HPA1HPA4*).

For the first incompatibility, in *HPA*, we found pseudo-heterozygous LoF variations including two premature stop-codons and three frameshifts at both reference copies of *HPA* (Fig. 4c and Supplementary Table 9) due to the repetitiveness of the gene sequences. As some accessions did not show expression of *HPA2* most likely due to hypermethylation of their promoters, we tested which of the *HPA* alleles was present in the RNA-seq read data to assign the LoF to either of the *HPA* copies (Fig. 4b). With this, we could assign one stop-codon gain (LoF-300) and two frameshifts (LoF-170, LoF-230) to *HPA2* because these LoF alleles were absent in the RNA-seq data in the accessions which lacked *HPA2* expression. Notably, the frameshift LoF-170 could be rescued by alternative splicing as observed in RNA-seq read mapping (Supplementary Fig. 4). Furthermore, cytosine methylation profiles revealed hypermethylated promoters of *HPA2* in 340 accessions and of *HPA1* in 50 accessions (Fig 4d, e), suggesting gene silencing in these accessions, which was in agreement with the absence of pseudo-heterozygous variants in the RNA-seq read data (Fig. 4b and Supplementary Fig. 4).

By combining the LoF variant genotyping and the methylation analyses, we found 767 accessions with a non-functional *HPA2* (*HPA2*^*−/−*^ *HPA1*^+*/*+^) allele and five accessions with a non-functional *HPA1* (*HPA2*^+*/*+^*HPA1*^*−/−*^) allele (Fig. 4f and Supplementary Table 10). Also, 49 accessions did not feature any functional alleles (*HPA2*^*−/−*^*HPA1*^*−/−*^) and were expected to carry additional modifier(s) to complement the loss of functional copies. To find the locations of these modifiers, we used the absence of functional copies as phenotype (Supplementary Data 3) to run a GWA under the mixed linear model using the SNP markers from the 1001 Genomes Project (www.1001genomes.org). This GWAS revealed a significantly associated region at chr4:11.00-11.15 Mb (Fig. 4g and Supplementary Fig. 5, alpha level of 0.05, Bonferroni correction), corresponding to the *HPA3* locus found in Kyo (chr4:11.11 Mb). Though other peaks in unlinked regions were present, these additional loci explained only a small proportion of the heritability.

**Figure 5:**
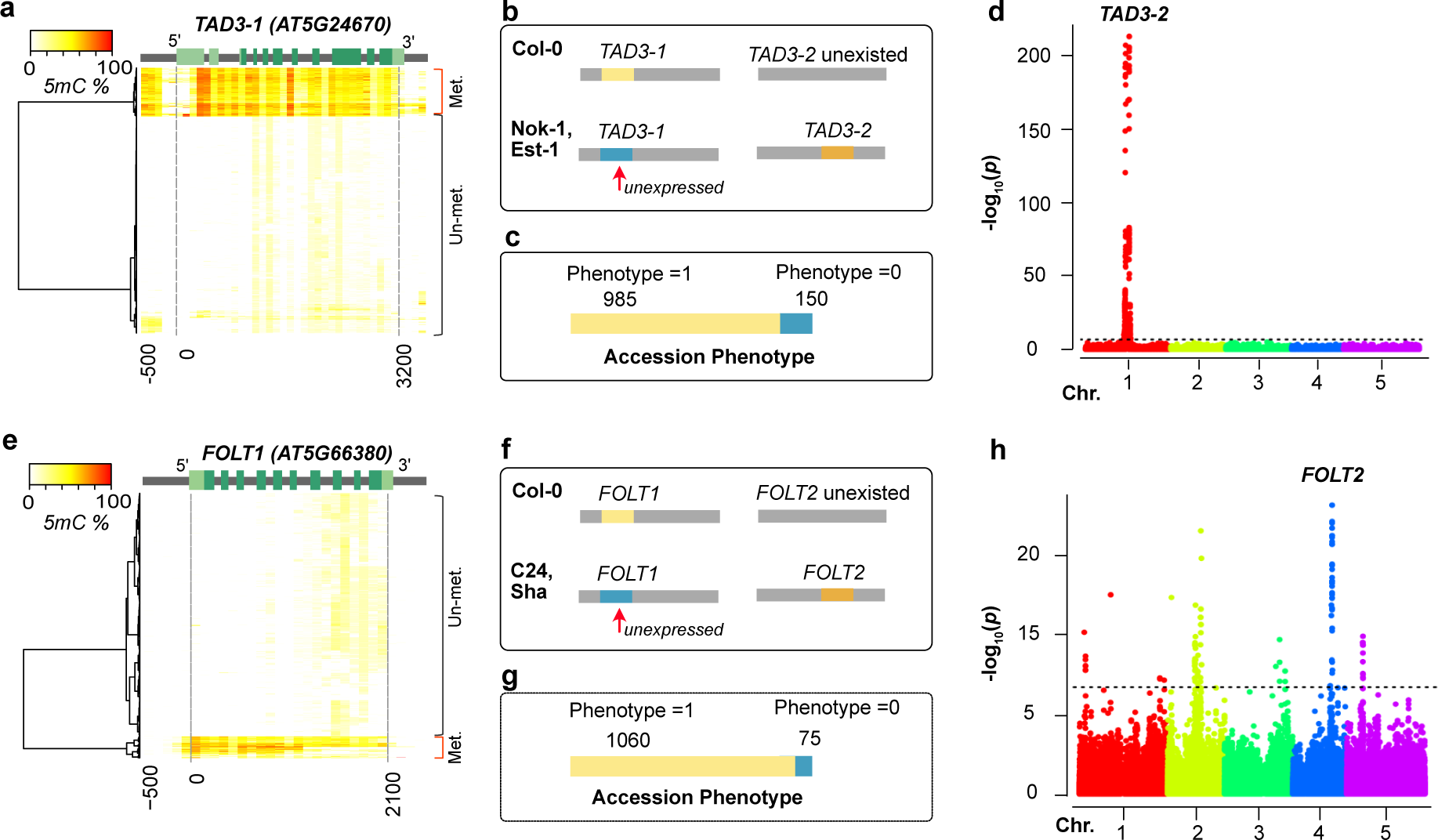
Mapping non-reference gene copies of incompatible alleles using GWAS. **(a, e)** Hierarchical clustering of cytosine methylation profiles in *TAD3-1* (a) and *FOLT1* (e) based on 888 *A. thaliana* accessions from 1001 Epigenomes Project (Kawakatsu et al. 2016). The methylation profile was calculated based on 100 bp sliding windows from the 500 bp upstream of transcription start site to the 300 bp downstream of the transcription end site. Light green: UTR, green: coding region, gray: intron or up/down-stream. **(b, f)** The genetic incompatibilities introduced by *TAD3* (b) in hybrids between Col-0 and Nok-1/Est-1, and by *FOLT* (f) in hybrids between Col-0 and C24/Sha as shown previously (Durand et al. 2012; Agorio et al. 2017). **(c, g)** The number of accessions with functional (Phenotype=1) or non-functional (Phenotype=0) *TAD3-1* (c) and *FOLT1* (g) gene copies. **(d, h)** Manhattan plots of the GWA using the absence of functional *TAD3-1* (d) or *FOLT1* (h) gene copies as phenotype. The dashed line indicates the significance threshold after multiple testing correction (−log_10_*P*=6.68).

Analyzing the haplotypes at the 39 significantly associated SNP markers at *HPA3* locus revealed a somewhat, but not entirely homogenous haplotype in many of the *HPA2*^*−/−*^ *HPA1*^−/−^ carriers (Fig. 4h). To confirm that the modifying haplotype is still identical to the Kyo allele, we aligned the short reads of all accessions of the 1001 Genomes Project against the Kyo reference sequence and found that overall 162 accessions carried the *HPA3* allele (Supplementary Data 3). However, unexpectedly the Kyo *HPA3* was only found in 13 of the 49 accessions without functional *HPA2* and *HPA1* alleles, suggesting the presence of two different modifiers at this locus on chromosome 4. To find more support for this, we ran a new GWA without the 162 *HPA3* (Kyo-like allele) carriers, which still led to a significantly associated locus at the region of *HPA3* (Supplementary Fig. 6). Further short-read mappings of all 1,135 genomes against the Cvi-0 reference sequence (where only one copy of *HPA* exists) revealed the presence of (at least) two additional copies (in addition to *HPA1* and *HPA2*) in a total of 42 accessions including all the 36 accessions with the unknown rescuing alleles (Fig. 4i and Supplementary Data 3). Together this suggested that *HPA3* also rescues incompatible *HPA1* and *HPA2* allele combinations in natural populations and that the locus of *HPA3* contains an additional haplotype (hereafter named *HPA4*) which also rescues the incompatibility between *HPA1* and *HPA2*.

**Figure 6.**
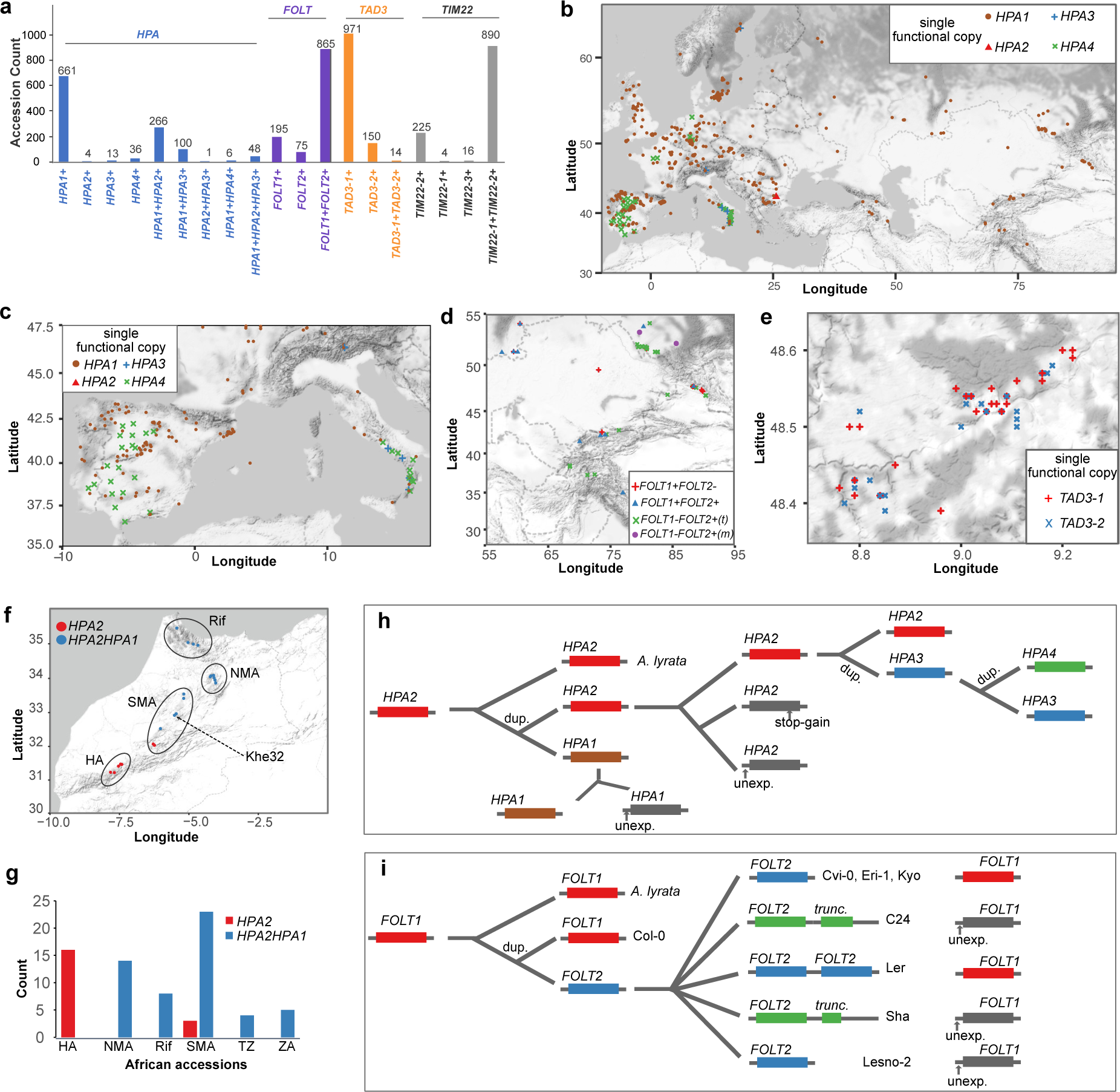
Distribution, origin and evolution of incompatible alleles. **(a)** The number of accessions with different functional copies of *HPA, FOLT, TAD3* and *TIM22* across 1,135 *A. thaliana* accessions from the 1001 Genomes Project. +: functional. **(b)** Geographic distribution of accessions only with one functional copy of *HPA*. **(c, d, e)** Examples of geographically close accessions with only one functional copy of *HPA* in Southern Europe (c), *FOLT* in Central Asia (d) and *TAD3* in Germany (e). *FOLT1*-*FOLT2*+(t): accessions with a functional *FOLT2*, a truncated *FOLT2* and an unexpressed *FOLT1. FOLT1*-*FOLT2*+(m): accessions with a functional *FOLT2* and an unexpressed *FOLT1*, but without truncated *FOLT2* copies. **(f, g)** Gene copies of *HPA* in African *A. thaliana* accessions. TZ: Tanzania, ZA: South Africa. **(h, i)** Schematic of a possible and parsimonious evolutionary history of gene duplication and non-functionalization of *HPA* (h) and *FOLT* (i). unexp.: unexpressed. stop-gain: premature stop codon gained SNP. dup.: gene duplication.

We continued to apply the same approach to the other three incompatibilities. For *TIM22*, 16 accessions of the 1001 Genome Project revealed non-functional allele combinations (Supplementary Table 11 and Supplementary Data 4), again indicating the existence of modifying alleles. However, a GWAS using the presumably-incompatible allele combinations as molecular phenotype did not reveal only one, but numerous significantly associated loci (Supplementary Fig. 7). This might be explained by the low number of incompatible allele carriers, which could affect the power of association mapping leading to false-positive associations. However, even though we could not locate the modifying allele, further analysis of read mapping coverage in *TIM22* revealed that all 16 accessions with non-functional *TIM22-1/TIM22-2* allele combinations carried at least one additional third copy of *TIM22*, while among all 1,119 other genomes of the 1001 Genomes Project only three genomes carried additional copies. This suggests that, like for *HPA*, known non-functional allele combinations of *TIM22* are in fact rescued by additional copies. Moreover, in ten of the 16 accessions, *TIM22-2* showed not only one but multiple additional copies (hereafter named as *TIM22-3*), which was further supported by the genome assembly of Ty-1 (https://genomevolution.org/CoGe/GenomeInfo.pl?gid=54584, unpublished), where we could find a cluster of four tandemly arranged *TIM22* gene copies at the *TIM22-2* locus.

Because the reference sequence only contained one copy of *TAD3 (TAD3-1)* (Agorio et al. 2017) and *FOLT (FOLT1)* (Durand et al. 2012), we modified our GWA method and only used non-functional alleles at the reference gene as the phenotype to map modifiers for the two remaining incompatibilities (Fig. 5 and Supplementary Fig. 8). Due to this modification we would expect to map also the location of the duplicated genes as we had found them in the AMPRIL founders. For *TAD3*, an essential ortholog of the yeast tRNA Adenosine deaminase 3 (Gerber and Keller 1999), we did not find any accessions with LoF alleles, but we found 150 accessions with a hypermethylated promoter in *TAD3-1* similar to the methylated promoters found in Nok-1 and Est-1, which are known to be non-functional due to this methylated promotor region (Agorio et al. 2017) (Fig. 5a, b). The GWA result revealed one significant peak (Fig. 5c,d), which as expected co-located with the *TAD3-2* locus (Agorio et al. 2017). All the 150 accessions carried multiple additional copies of *TAD3-2* (Supplementary Data 5), similar to Nok-1 and Est-1. This suggests that the expression silencing of *TAD3-1* is common in natural populations and that the rescue of this loss-of-function allele is generally mediated by additional gene copies at the *TAD3-2* locus as it was shown in the original description of this incompatibility (Agorio et al. 2017).

The last of the four genetic incompatibilities, introduced by *FOLT* encoding for a folate transporter, was previously discovered in hybrids from crosses between Col-0 x C24/Sha (Törjék et al. 2006; Simon et al. 2008; Durand et al. 2012). Col-0 has only one copy of *FOLT* (*FOLT1*) at chromosome 5, whereas the C24 and Sha have an additional copy, *FOLT2*, at chromosome 4 including some extra truncated copies near *FOLT2* (Supplementary Table 12). In the earlier study, the truncated copies were shown to express siRNAs and activate the RNA-directed DNA methylation pathway to silence *FOLT1* in C24 and Sha (Durand et al. 2012), which resulted in a lethal allele combination in F2 hybrids between Col-0 (*FOLT1*^+*/*+^) and C24/Sha (*FOLT1*^*−/−*^ *FOLT2*^+*/*+^) (Fig. 5f). Analyzing the accessions of the 1001 Genome Project for functional and non-functional alleles of *FOLT*, we found 75 accessions with methylated promoters of *FOLT1* likely leading to expression silencing (Fig. 5e), which was supported by the lack of *FOLT1-*specific pseudo-heterozygous SNPs in the RNA-seq data (Supplementary Fig. 9). Besides evidencing the expression silencing of *FOLT1*, this also suggested the existence of additional *FOLT* gene copies in these 75 accessions. When we repeated our GWA approach to find these interacting loci of *FOLT1*, we found multiple significantly associated loci including one region corresponding to *FOLT2* (Fig. 5g, h).

Further analyses of the *FOLT* gene copies using short read mapping against the C24 and Sha reference sequences revealed the presence of *FOLT2* in all the 75 accessions with methylated promoter of *FOLT1* and truncated copies of *FOLT2* genes in only 46 out of such 75 accessions (Fig. 5e, Supplementary Table 13 and Supplementary Data 6) suggesting that the methylation of the *FOLT1* promoter remains stable even after the truncated copies are segregated out. This is coherent to what was shown in successive generations of Sha x Col-0 RILs where the inducing locus was segregated away six generations ago (Durand et al. 2012).

Taken together we found evidence that all four incompatible allele combinations, initially identified within artificial intercross population, also occur in nature, and that the incompatible allele combinations of three of them were surprisingly common. For all incompatible allele combination carriers, we found evidence for the existence of additional copies, and could even map the locations of some of those using GWA.

### Geographic distribution of incompatible allele combinations

We next asked how prevalent the potential for incompatible allele combinations was within the natural population of *A. thaliana* by analyzing the presence of different haplotypes in different geographic regions. For this, we first analyzed the allele frequencies of different haplotypes (i.e. the different combinations of functional alleles within individual plants), which varied substantially across the accessions of the 1001 Genomes Project (Fig. 6).

For example, among the accessions that carried only one functional copy of *HPA*, we found that 661, 4, 13 and 36 accessions with a single functional copy of *HPA1, HPA2, HPA3*, or *HPA4*, respectively (Fig. 6a, b, Supplementary Fig. 10). Similarly, most accessions only featured one functional copy of *TAD3* including 971 accessions with only *TAD3-1* and 150 with only *TAD3-2* (Fig. 6a, Supplementary Table 14 and Supplementary Fig. 11). In contrast, most of the accessions included multiple functional alleles of *FOLT* as well as of *TIM22* (Fig. 6a).

Genetic incompatibilities become effective in the offspring of accessions with incompatible alleles. In particular the offspring of accessions with only one functional gene copy will lead to the highest reduction in fitness. We found numerous accessions with the potential to create incompatible allele combinations in their offspring in geographically close regions (Fig. 6c, d, e). For example, among 30 accessions in the South of Italy we found 9 accessions with only a functional *HPA1* closely located with 14 accessions with either a functional *HPA3* or a functional *HPA4* only (Fig. 6c). Likewise, accessions with contrasting functional copies of *FOLT* and *TAD3* were collected from the same regions in Central Asia and Germany (Fig. 6d and e) evidencing that incompatible alleles could segregate within the same populations.

Interestingly, within some of these hybrid zones of accessions with incompatible alleles, we also found accessions with multiple functional gene copies. For example, in South Italy there were 7 of the 30 accessions that featured multiple functional copies of *HPA*, and similarly in Central Asia, we could find multiple accessions with two functional copies of *FOLT* in addition to the accessions with only one functional copy (Fig. 6d and Supplementary Fig. 10-12). In contrast, however, we only observed a few accessions with multiple functional alleles in the hybrid zone of incompatible *TAD3* alleles, most likely because only 14 such accessions were found in the entire set of accessions. Even though we could not find evidence that such hybrid zones are enriched for additional copies or that those additional copies would evolve in these regions, the presence of haplotypes with additional gene copies in these hybrid zones have the potential to mediate the gene flow between the haplotypes with the incompatible allele combinations.

### Origin and evolution of incompatible alleles

To figure out when these incompatible alleles originated and how they evolved, we investigated their ancestral genotypes within African genotypes which likely represent the ancestral populations of the Eurasian accessions (Durvasula et al. 2017).

While almost all (99.7%) of the Eurasian accessions had both *HPA2* and *HPA1* (either functional or not), we found that only 27% (20 out of 75) of the African accessions carried only *HPA2* (Fig. 6f, g and Supplementary Data 3). This suggested that *HPA2* was the ancestral copy of *HPA* which was further supported by the synteny alignment with the close relative *Arabidopsis lyrata* (Blevins et al. 2017), which only featured a single *HPA* gene in a region syntenic to *HPA2* and that the duplication events leading to *HPA1* happened early on in Africa (Fig. 6h). One accession, Khe32, from Morocco carried *HPA1* along with the most frequent LoF variant LoF-300 in *HPA2* in the Eurasian accessions, suggesting that the first accession with only a functional *HPA1* possibly arose in North-West Africa and thereby suggests that the genetic incompatibility with *HPA2* carriers was already possible before the Eurasia colonization of *A. thaliana*.

In contrast, *HPA3* and *HPA4* only occur in Eurasian accessions, most likely indicating that the additional duplication events happened later (Fig. 6h). Comparing the gene sequences of the *HPA* genes revealed that *HPA3* was duplicated from *HPA2* (Supplementary Fig. 13), while *HPA4* might be a tandem duplicate of *HPA3* as it is likely located close to *HPA3*. Interestingly, *HPA3* and *HPA4* carriers segregated for the ancestral LoF-300 variant at *HPA2* (*HPA3*: 68 with and 94 without; *HPA4*: 10 with and 32 without, Supplementary Fig. 14), suggesting free segregation of different *HPA* alleles. In striking contrast to this, the inactive alleles of *HPA1* (i.e. alleles with a non-functional *HPA1* sequence, but excluding the accessions with full deletion alleles as found in Cvi-0) were almost perfectly coupled (48 of 50) with inactive alleles of *HPA2*. Such carriers nearly all had *HPA4* (35) or *HPA3* (13) (Supplementary Fig. 15) and were mainly located in the Iberian Peninsula and Southern Italy suggesting that the additional *HPA* copies were necessary to buffer the incompatibility and to allow the foundation of these populations (Fig. 6c).

Unlike the incompatibility in *HPA*, genetic incompatibilities can also arise in recent population history. All African accessions only had the *TAD3-1* copy, while accessions with multiple copies of *TAD3* can only be observed in Eurasia (Supplementary Fig. 10). Similarly, 38 of the African accessions only have *FOLT1*, while the other 37 accessions have both *FOLT1* and *FOLT2*, but none of the accessions featured a truncated copy of *FOLT2* (*FOLT2tr*), which is the mechanistic origin of the incompatible alleles at *FOLT1* and *FOLT2* (Supplementary Fig. 16 and Supplementary Data 6). In contrast, within the Eurasian accession, we even found three different haplotypes of *FOLT2tr* within a total of 46 accessions based on short read alignments against the eight *A. thaliana* genomes (Jiao and Schneeberger 2020) (Supplementary Fig. 17 and Supplementary Data 6). This together suggests that also the genetic incompatibility that is based on *FOLT2tr* has evolved recently after the migration *A. thaliana* to Eurasia (Fig. 6i).

## Discussion

Although gene duplication provides genetic backup of essential genes, duplicated genes can also lead to incompatible allele combinations when the duplicated genes undergo reciprocal pseudo-functionalization in separate genomes. Here we studied incompatible allele combinations of four duplicated gene pairs by integrating genetic and genomic information using a multi-parental intercross population leading to the identification of four genetic incompatibilities.

Unexpectedly in this population we identified some lines with what initially looked like an incompatible allele combination of one of the incompatibilities, as they could be rescued by an additional, so-far unknown gene copy. Encouraged by this, we developed a GWA method to map modifying alleles also in natural populations by integrating genomic and epigenomic data to generate a molecular phenotype that describes potentially incompatible allele combinations. With this we identified many natural accessions with putatively incompatible allele combinations and could elucidate the genetics of these incompatibilities as they occur in natural population using the 1,135 accessions released by the 1001 Genomes Project (Alonso-Blanco et al. 2016). This implies that besides presence/absence analysis as performed by standard pan-genome analysis, gene location is an essential feature of a gene (or gene family).

Based on multi-omics data from hundreds of *A. thaliana* accessions, we could comprehensively describe the origin and evolution of several incompatible allele combinations of the four incompatibilities. We found that incompatible alleles are surprisingly frequent in nature and also occur in sympatry, suggesting that the evolution of genetic incompatibilities does not require separated populations as proposed by BDM model (Bateson 1909; Dobzhansky 1937; Muller 1942). Moreover, such incompatible allele combinations can persist over long periods as some of the alleles studied here may have originated before *A. thaliana* colonized Eurasia.

Here, new gene copies arose either from distal duplications (in *HPA* and *FOLT*) or tandem duplications (in all four cases) and counteract incompatibilities. Even though additional copies reduce the frequency and impact of genetic incompatibilities, they could also increase the potential for more incompatible allele combinations. Subsequent sub-functionalization could be a way out of the trap of genetic incompatibilities, as there would be a selective pressure to keep both gene copies. While we have assumed functional redundancy of all gene copies in this work, an extensive number of accessions shared the functional copies of bot *FOLT* genes (865 of 1,135), which could indicate that these copies are not fully redundant and thereby limit the establishment of incompatible allele combinations.

Taken together, our work demonstrates that the potential for genetic incompatibilities due to duplicated essential genes is surprisingly high in nature. However, the effects of such incompatibilities are counteracted by additional gene copies, which undergo dynamic changes shaped by the recurrent events of gene duplication and non-functionalization during population history.

## Materials and Methods

### AMPRIL construction

The eight *A. thaliana* accessions An-1, C24, Col-0, Cvi-0, Eri-1, Kyo, L*er*, and Sha were selected as the AMPRIL founders. We previously constructed a first version of AMPRIL population (AMPRIL I) including six RIL subpopulations (Huang et al. 2011). Here AMPRIL I was extended with six additional subpopulations (called AMPRIL II) based on different pairwise intercrosses of the eight founders (Fig. 1a). Each subpopulation contains approximately 90 individuals (Supplementary Data 1). All plants were grown under the normal growth conditions in greenhouse at the MPI-PZ (Huang et al. 2011). DNA of 1,100 samples was extracted from the flower buds and prepared for RAD-seq sequencing (Baird et al. 2008).

### RAD-seq Library preparation and sequencing

All plants were grown in the greenhouse. The total DNA from each of the 1,100 samples was extracted from the flower buds using DNeasy Plant Kit 96 (Qiagen) and eluted in 200 µl Elution buffer (EB). DNA of each genotype was isolated twice. Both genotype samples were pooled in 1.5 ml tube and the DNA was concentrated by isopropanol precipitation for 2h at −20°C. The samples were centrifuged at 12000 *g* for 5 min at 4°C, the supernatant was removed and the DNA pellet was washed with ice-cold 70% ethanol. Centrifugation was repeated, the supernatant was removed. Air-dried DNA was resuspended in a nuclease free water to 26 ng/µl and stored at −20°C until use. RAD-seq sequencing libraries were prepared as described (Etter et al. 2011) with modifications. Per genotype, 500 ng DNA were digested with 10 units of CviQI (NEB, cutting site G’TAC) at 25°C for 2 h. The number of expected cutting sites was estimated around 236,000 based on the Col-0 genome sequence. Cut DNA was purified using 96 DNA clean and concentrator kit (Zymo) a diluted in 25 µl EB. The 192 different (selected out of total 210 designed, Supplementary Data 1) P1 adapters (200 nM) containing unique 12 bp barcodes were ligated by incubation with T4 ligase (NEB) at room temperature for 30 min and the reaction was terminated by 20 min at 65°C. After 30 min at room temperature, 5 µl from each 192 P1-barcoded sample were combined in a 2 ml low bind tube. 3 x 130 µl aliquots were transferred to fresh tubes and DNA was fragmented to average size of 500 bp using Covaris. Sheared DNA was purified using QiaMinElute columns (Qiagen), eluted in 10 µl EB, and the three samples were first pooled and then divided into two 15 µl samples that were run on 1% agarose gel. Regions of 300-500 bp fragments were dissected and DNA was isolated using MinElute gel extraction kit (Quiagen), eluted in 10 µl EB, the samples were pooled, DNA fragment ends were repaired using Quick Blunting™ kit (NEB), purified with QIAquick column (Qiagen) and eluted in 43 µl EB. 3’ deoxy-adenine overhangs were added using Klenow Fragment (NEB), the sample was purified with QIAquick column, eluted in 45 µl EB, the P2 adapter was ligated, the sample was purified with QIAquick column and eluted in 53 µl EB. To determine the library quality, 10 µl RAD library were PCR amplified (1: 98°C 30 sec; 2: 14x 98°C 10 sec, 67°C 30 sec, 68°C 30 sec; 3: 68°C 5min, 4: 4°C hold) using NEB Next High-Fidelity master mix (NEB) in 25 µl reaction volume using RAD-Marker for/ RAD-Marker rev primers (25 nM each) and 5 µl PCR product were loaded into 1% agarose gel next to the 1 µl RAD library template. If the PCR product smear was at least twice as intense as the template smear, the library was considered as of high quality and the amplification was repeated in 50 µl reaction volume. The product was cleaned using AMPure magnetic beads (Beckman Coulter) and dissolved in 20 µl EB. Finally, the sample was run on 1% agarose gel, the region of 300-500 bp fragments was cut out, DNA was isolated using MinElute Gel extraction kit (Qiagen) and eluted in 20 µl EB. The library was sequenced in an Illumina Hiseq2000 sequencing machine. Sequencing reads were demultiplexed according to the barcodes (Supplementary Data 1).

### AMPRIL genotyping

We used previously released whole-genome short read data(Jiao and Schneeberger 2020) to generate markers for genotyping the AMPRILs. We mapped the reads to the reference sequence and called SNPs using SHORE (version 0.9) (Ossowski et al. 2008) with default parameter settings. Only homozygous SNP calls were selected after removing SNP calls with low quality (quality < 30), in the repetitive regions or in regions with low mapping quality (quality < 30). The actual marker sets for each of the twelve subpopulations (ABBA, ACCA … GHHG) were selected based on respective parental genomes, exclusively selecting bi-allelic SNP markers. After excluding samples with replicates or too few sequencing reads, we performed genotyping on 992 AMPRILs using a Hidden Markov Model-based approach similar to the recently presented method for the reconstruction of genotypes derived from two parental genomes (Rowan et al. 2015) (for a detailed description see Supplementary Note 1).

### Identifying genetic incompatibilities based on duplicated genes

Duplicated gene pairs were selected based on gene family clustering of protein-coding genes from all eight parental genomes using OrthoFinder (version 2.2.6) (Emms and Kelly 2015). We only selected inter-chromosome duplicated genes to avoid the effects of intra-chromosome linkage (Supplementary Data 2). For each duplicated gene pair, we required two copies in the reference sequence and at least one copy in one of the other genomes (DupGene2), or one copy in reference sequence and at least one copy on a different chromosome in at least one of the other parental genomes (DupGene1). We assessed the genotypes of each of the gene copies in each AMPRIL using the genotypes predicted in the middle of the respective reference gene, or in the case a gene was not present in the reference sequence – using the midpoint between the two closest flanking syntenic regions of the non-reference gene copy (based on synteny calculations from a previous study (Jiao and Schneeberger 2020)).

We predicted candidate genetic incompatibilities using a two-steps approach. In the first step, we performed chi-square tests (Equation 1) to check whether the frequency of allele pairs in duplicated genes was significantly distorted in any of the subpopulations or in any of two merged subpopulations (ABBA and EFFE or CDDC and GHHG which shared the same four founders, respectively).

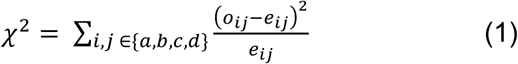

Here, the *o*_*ij*_ and *e*_*ij*_ represent the observed and expected allele pair frequency of duplicated genes, respectively, and *a, b, c, d* represent the parental genotypes in each subpopulation.

Additionally, we applied a modified chi-square test (Equation 2) for all the duplicated genes in the whole AMPRIL population by considering the effects of population structure.

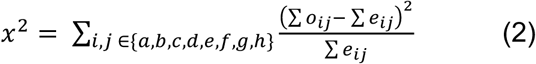

Here, the *o*_*ij*_ and *e*_*ij*_ represent the observed and expected allele pair frequency of inter-chromosome duplicated genes, respectively, and *a, b, c, d, e, f, g*, and *h* represent all parental genotypes. The observed and expected allele pair frequency in the whole population was the sum of observed (∑ *o*_*ij*_) and expected (∑ *e*_*ij*_) allele pair frequencies in each of the subpopulations.

The gene pairs with at least one significant segregation distortion in their observed allele combinations (FDR < 0.05) were kept for the next step. In this second step, we checked whether the respective gene copies contained loss-of-function variation or methylated promoters (as described below). To address alternative splicing which is known to rescue loss of-function variation, we also checked the gene annotation within the parental genome assemblies to confirm the loss-of-function. Only duplicated genes with confirmed loss-of-function alleles in both copies of the duplication (in at least one of the parental genomes) were kept as candidates for genetic incompatibilities based on duplicated genes.

### Identification of LoF variants in candidate genes

We mapped all whole-genome resequencing reads of 1,135 accessions from the 1001 Genomes Project (Alonso-Blanco et al. 2016), 75 accessions from Africa (Durvasula et al. 2017) and 118 accessions from China (Zou et al. 2017), to the Col-0 reference genome (TAIR10) (The Arabidopsis Genome Initiative 2000; Lamesch et al. 2012) using BWA (version 0.7.15) (Li and Durbin 2009) with the default parameter settings. SNPs and small indels were called using SAMtools (version 1.9) using default parameters (Li et al. 2009). Homozygous variants with mapping quality of more than 20 and with at least four reads aligned were kept. Pseudo-heterozygous variants in *HPA* and *TIM22* were also recorded. The large insertion and deletions in the 40kb extended genic region of focal genes were predicted using Pindel (version 0.2.5) (Ye et al. 2009) with parameter settings “-T 1 -x 5 -k -r -j” and Delly (version 0.8.1) (Rausch et al. 2012) with parameter settings “delly call -q 20 -r 20 -n -u 20 -g”. The functional effects of these variations were annotated using SnpEff (version 4.3p) (Cingolani et al. 2012) using the default parameter settings. The loss-of-function (LoF) effects include loss of start codon, loss of stop codon, gain of premature stop codon, damage of splicing acceptor or donor sites, frameshift and CDS loss.

### Clustering of cytosine methylation profile in gene promoter

Cytosine methylation data (the tab separated file of methylated cytosine positions) of 1,211 samples from the 1001 Epigenomes project (Kawakatsu et al. 2016) were downloaded from NCBI (927 samples under GEO accession GSE43857 and 284 samples under GEO accession GSE54292). After removing the redundant data sets 888 and 161 data sets from GSE43857 and GSE54292, respectively, were retained. For each sample, we calculated the percentage of methylated cytosines in CG, CHG and CHH contexts from 500bp upstream of the transcription start sites to 300bp downstream of the transcription termination sites of each candidate gene in 100bp non-overlapping sliding windows. These methylation profiles were hierarchically clustered using the hclust function implemented in R (version 3.5.1). The pairwise, Euclidean distances between all methylation profiles were calculated and Ward’s method was used to cluster the samples into two groups (hypermethylated and unmethylated). This clustering was performed for the samples of GSE43857 and GSE54292 separately as these two data sets were processed in two studies with different pipelines (Dubin et al. 2015; Kawakatsu et al. 2016). The heatmap of methylation patterns was drawn in R using the heatmap.2 function.

### Analysis of DNA methylation within the genomes of the AMPRIL founders

For six accessions (Col-0, An-1, C24, Cvi-0, Ler, Kyo), we downloaded the whole-genome DNA methylation data from NCBI from the 1001 Epigenomes Project (Kawakatsu et al. 2016) (GSE43857). For Eri-1 and Sha, DNA methylation data was generated using whole-genome bisulfite sequencing by the Max Planck Genome center. DNA was extracted from plants grown in the greenhouse under standard conditions using the Qiagen DNEasy Plant Mini Kit (Qiagen, Germany) and a sequencing library was prepared using the NEXTflex Bisulfite Library Prep Kit. This library was sequenced on an Illumina HiSeq2000 machine. Sequencing reads were aligned the reference sequence using Bismark (version 0.20.0) (Krueger and Andrews 2011) with these parameters “-q --bowtie2 -N 1 -L 24 -p 20”. The cytosine methylation profiles in candidate genes were calculated using the same sliding window method as described above. The cytosine methylation profiles together with the profiles from GSE43857 were then clustered again with the same clustering method as described above.

### Mapping modifiers of incompatible alleles using GWA

We predicted the presence and absence of functional copies of duplicated genes (*HPA, TIM22*) in each of 1,135 accessions from the 1001 Genomes Project according to the annotations of loss-of-function variations and the clustering of cytosine methylation profiles in promoters (Supplementary Data 3-6). For accessions without available methylation sequencing data, we assume the focal genes are expressed. The presence or absence of any functional copies of the reference genes were used as binary phenotype (presence: 1, absence: 0). For the association, we selected 238,166 high-quality SNP markers (minor allele frequency >0.05 and missing rate < 0.1) from 1001 Genomes Project and imputed missing alleles with IMPUTE2 (Howie et al. 2009; Howie et al. 2012). An in-house R script implementing the mixed linear model with correction of kinship bias was used to perform the GWA (Bonferroni correction, P < 0.05).

### Copy number variation analysis

To test for the existence of *HPA3* in a genome, we mapped whole-genome short reads of 1,135 accessions from 1001 Genomes Project (Alonso-Blanco et al. 2016), of 75 accessions from Africa (Durvasula et al. 2017) and of 118 accessions from China (Zou et al. 2017) to the Kyo genome assembly (Jiao and Schneeberger 2020) using BWA (version 0.7.15) with default parameters. Accessions with an average mapping coverage >= 5 along the *HPA3* region and duplicated region breakpoints (identified based on the sequence alignment against Col-0 genome using SyRI (Goel et al. 2019) (version 1.0) with default parameters) were considered as *HPA3* carriers.

The copy number of duplicated genes was estimated by the ratio between the average mapping coverage within the focal gene and the average mapping coverage across the whole genome. To estimate the copy number of *HPA*, we mapped the short reads to the Cvi-0 genome assembly using BWA (version 0.7.15) with default parameters as the Cvi-0 genome only has one copy of *HPA*. For *TAD3, FOLT* and *TIM22*, the copy number was predicted based on short reads mapping against the reference genome where both *TAD3* and *FOLT* only have one copy. For the *TIM22*, copy number was estimated based on the average mapping coverage at both *TIM22-1* and *TIM22-2*.

### Fine-mapping of incompatible allele at LD2 in Cvi-0 x Col-0 RIL

After the observation that it was not possible to obtain a RIL homozygous for the Col-0 allele at LD2.1 while homozygous for the Cvi-0 allele at LD2.3 (Simon et al. 2008), we derived two distinct heterogeneous inbred families (HIF) from two segregating RILs from this population (Supplementary Fig. 2). 8RV467 is segregating for a region largely encompassing LD2.1 while fixed Cvi-0 at LD2.3. 8RV408 is segregating for LD2.3 while fixed Col-0 at LD2.1. In each derived HIF family, it is again not possible to fix the incompatible allele combination, so the families were used to exclude intervals that do not contribute to the incompatible interaction. Two rounds of fine-mapping were conducted by genotypically screening increasing populations of descendants (at the seedling stage) for recombinants in the interval. Gradually, the causal interval is reduced and delineated by markers exploiting known SNPs and indels in the Cvi-0 sequence.

### Segregation of T-DNA mutant line at gene *TIM22-1* and *TIM22-2*

T-DNA lines GABI_848H04 segregates for an insertion in *TIM22-1*, while GABI_626H10 segregates for an insertion in *TIM22-2*, both in a Col-0 background. Descendance were screened genotypically at the seedling stage to characterize the segregation of the T-DNA insertion allele.

### Complementation cross to validate the incompatible alleles of *TIM22*

A LD2.3 HIF line (8HV408-Het) was crossed with a GABI_626H10 line, both at the heterozygous state, in order to segregate potentially for all four possible hybrid allelic combinations at LD2.3, while maintaining a fixed Col-0 allele at LD2.1. The F1 descendance (123 individuals, from both cross directions) was screened genotypically at the seedling stage for the presence/absence of both the T-DNA insertion and the Cvi-0 allele.

## Data availability

Raw RAD-seq data of the AMPRIL population was deposited to European Nucleotide Archive (ENA) under the project accession ID PRJEB39883. Genome resequencing data of all eight founders and BS-seq data of Eri-1 and Sha can be accessed in ENA under the project accession ID PRJEB31147 and PRJEB38624. Genome resequencing data generated in 1001 Genomes Project (Alonso-Blanco et al. 2016) was downloaded from NCBI under the project ID SRP056687. RNA-seq and methylation data from the 1001 Epigenomes Project (Kawakatsu et al. 2016) were downloaded from NCBI under the project accession ID GSE80744, GSE54680, GSE43857 and GSE54292. The SNP markers from the 1001 Genomes Project were downloaded from https://1001genomes.org/data/GMI-MPI/releases/v3.1/.

## Code availability

Custom code used in this study can be freely accessed at https://github.com/schneebergerlab/AMPRIL-GI

## Seed availability

We are currently preparing seeds of new AMPRIL populations for submission to NASC. (http://arabidopsis.info/). Note, heterozygous regions as reported in the genotype data might be fixed for one of the alleles in the requested seeds.

## Acknowledgements

The authors would like to thank Detlef Weigel (Max Planck Institute for Developmental Biology, MPI-EB), Felix Bemm (MPI-EB), Todd Michael (Salk Institute), Joe Ecker (Salk Institute), Florian Jupe (Salk Institute) for releasing Ty-1 assembly prior to publication, Mehmet Goktay (Max Planck Institute for Plant Breeding Research, MPI-PZ) for help with the interpretation of genetic variation in Africa, Pádraic J. Flood (MPI-PZ) for helpful comments on the manuscript, the Max Planck Genome center for their sequencing efforts and the 1001 Genomes (1001genomes.org) and Epigenomes consortia for releasing their data. This work was funded by the Deutsche Forschungsgemeinschaft (DFG, German Research Foundation) under Germany’s Excellence Strategy – EXC 2048/1– 390686111 (K.S.), and the European Research Council (ERC) Grant “INTERACT” (802629) (K.S.).

## Authors contributions

WBJ and KS designed this project. WBJ, VP, JK, and FL analyzed the data. PP and AP generated the RAD-seq data. OL, MF, IG and CC performed the mapping and complementation experiments of LD2/*TIM22*. SE and MK generated the AMPRIL population. WBJ and KS wrote the paper.

