## Supplementary Notes, Tables and Figures for "The evolutionary dynamics of genetic incompatibilities introduced by duplicated genes in *Arabidopsis thaliana*" (from another version of this preprint: version 1, 2020-09-22)

**Supplementary Data 1**. Summary of RAD-seq data of AMPRIL population

**Supplementary Data 2**. List of SNP markers used for genotyping AMPRIL population

**Supplementary Data 3.** Identity by descent (IBD) matrix. The eight characters from a-f depict the eight founder haplotypes. The first three columns show the coordinate of IBD haplotype blocks. The flowing eight columns show these parental haplotypes. The remaining 992 columns indicate the recombinants of parental haplotypes for 992 RILs

**Supplementary Data 4.** List of inter-chromosome duplicated genes.

**Supplementary Data 5.** Loss-of-function variation, promoter methylation profile clustering and copy number of *HPA* across 1,135 *A. thaliana* accessions from the 1001 Genomes Project, 75 African accessions and 118 Chinese accessions.

**Supplementary Data 6.** Loss-of-function variation, promoter methylation profile clustering and copy number of *TIM22* across 1,135 *A. thaliana* accessions from the 1001 Genomes Project, 75 African accessions and 118 Chinese accessions.

**Supplementary Data 7.** Loss-of-function variation, promoter methylation profile clustering and copy number of *TAD3* across 1,135 *A. thaliana* accessions from the 1001 Genomes Project, 75 African accessions and 118 Chinese accessions.

**Supplementary Data 8.** Loss-of-function variation, promoter methylation profile clustering and copy number of *FOLT* across 1,135 *A. thaliana* accessions from the 1001 Genomes Project, 75 African accessions and 118 Chinese accessions.

**Supplementary Note 1**

***Pre-genotyping***

The pre-genotyping was performed in each subpopulation separately. For each sub-population, SNP markers were selected based on the SNP markers identification between the respective parental genomes (Supplementary Data 2). Each sample was pre-genotyped at each of the SNP marker positions based on the number of aligned short reads under the equation 1 (Eq. 1). The probabilities of two homozygous paternal genotypes ($x_1$, $x_2$) and heterozygous genotype $f(a, b)$ were estimated as

$$x_1 = \frac{(a+b)!}{a! * b!} * 0.99^a * 0.01^b$$

$$x_2 = \frac{(a+b)!}{a! * b!} * 0.99^b * 0.01^a$$

$$f(a,b) = \frac{(a+b)!}{a! * b!} * p^a * p^b,\ p = 0.5 \qquad \text{(Eq.1)}$$

Here, the *a* and *b* represent the read counts for the reference and alternative allele. 0.01 is the assumed error rate for an allele as observed in the short-read alignments. This step of pre-genotyping was implemented in our previously released tool TIGER[1].

***Two stage Hidden-Markov-Models***

To translate the raw genotypes to identity-by-decent haplotype assignments, two different HMMs were developed for the homozygous (HMM-1) and the heterozygous (HMM-2) part of the genome. In HMM-1, five hidden states were defined and labelled as A, B, C, D (which represent the four parents of a subpopulation) and Z (which represents heterozygous regions). The emission states represent the possible observation values, which are from the pre-genotyping step. Regions labeled as Z were further analyzed by the HMM-2. The HMM-2 contains ten hidden states, where each hidden state has ten emission states. The ten hidden states represent all possible allele combinations (four homozygous and six heterozygous genotypes) from the offspring of hybrids of four parents. To reduce the errors of genotyping, the HMM-2 only considered SNPs markers, which were unique for one of the parents at that marker position. Besides, to reduce errors at the transitions between heterozygous and homozygous regions, in addition to the markers in the Z regions, 100 markers within the flanking homozygous regions were considered.

***Simulations and training of the HMM***

To train and validate our method, we firstly simulated a multiparent population with the same crossing scheme as the AMPRIL subpopulations including 5,000 samples. Three different average sequencing coverage rates (0.1x, 1x, and 10x) were simulated and 1.2 million markers were generated. The recombination rate was simulated using an extended version of the tool Pop-seq[2,3] (http://sourceforge.net/projects/popseq). Firstly, the probability matrix for the transition and emission was estimated using a supervised learning strategy by selecting 1,000 randomly simulated samples with an average sequencing coverage rate of 1x. Then, we applied the resulting HMM to ten randomly sequenced samples and changed slightly the emission probabilities by adding or removing values in the range from 0.01 – 0.001 until the genotype prediction was nearly correct with respect to the short-read data. For the genotyping results based on simulation data sets, we compared the predicted and expected number of crossovers. These comparisons suggested that the error rate itself was quite low, below 0.02% for 1x coverage rate.

To avoid over fitting due to the small testing data set, we ran the new model on all samples again and randomly selected ten new samples. For the new ten samples, we again compared the genotype prediction with the support from the short-read data. If the prediction was overall not correct, the emission probabilities were further stepwise changed and a new model was applied to the total sample set again. The procedure was performed until further changing of the emission values resulted in worse genotype predictions. To check the potential over-fitting, the HMMs were applied to a 1,000 new randomly simulated samples with 1x average sequencing coverage. To validate the prediction of our model, we compared the predicted genotypes of AMPRIL data with the previous genotyping result based on 300 SNP markers. For the HMM-2, the probabilities were trained based on the simulation population used for training the HMM-1 but only selecting regions which were heterozygosity. The emission probabilities were adjusted and validated as described above.

**Supplementary Table 1.** Loss-of-function (LoF) variation, expression (Exp.) and promoter methylation (Met.) of *TAD3* in the AMPRIL parental genomes.

| Acc. | TAD3-1 ID | TAD3-1 LoF | TAD3-1 Exp. | TAD3-1 Met. | TAD3-2 |
|---|---|---|---|---|---|
| An-1 | ATAN-5G34920 | - | + | - | - |
| C24 | ATC24-5G34790 | - | + | - | - |
| Col-0 | AT5G24670 | - | + | - | - |
| Cvi-0 | ATCVI-5G35090 | - | + | - | - |
| Eri-1 | ATERI-5G35050 | - | No RNA-seq data | - | - |
| Kyo | ATKYO-5G34660 | - | No RNA-seq data | + | + |
| L*er* | ATLER-5G34950 | - | + | - | - |
| Sha | ATSHA-5G34870 | - | + | - | - |

**Supplementary Table 2.** The observed and expected frequencies of allele pair combinations of *TIM22-1* and *TIM22-2* in the complete AMPRIL population.

| | ***Observed*** | ***TIM22-1*** | | | | | | | |
|---|---|---|---|---|---|---|---|---|---|
| | | An-1 | C24 | Col-0 | Cvi-0 | Eri-1 | Kyo | L*er* | Sha |
| ***TIM22-2*** | An-1 | 63 | 41 | 33 | 28 | 52 | 22 | 58 | 26 |
| | C24 | 13 | 98 | 12 | 17 | 46 | 23 | 36 | 21 |
| | Col-0 | 25 | 18 | 61 | 45 | 27 | 39 | 30 | 22 |
| | Cvi-0 | 5 | 13 | 4 | 66 | 12 | 1 | 4 | 18 |
| | Eri-1 | 38 | 35 | 18 | 23 | 61 | 23 | 18 | 18 |
| | Kyo | 21 | 27 | 40 | 25 | 21 | 70 | 20 | 35 |
| | L*er* | 25 | 54 | 18 | 19 | 19 | 17 | 71 | 8 |
| | Sha | 21 | 29 | 16 | 59 | 17 | 58 | 29 | 52 |
| | ***Expected*** | ***TIM22-1*** | | | | | | | |
| | | An-1 | C24 | Col-0 | Cvi-0 | Eri-1 | Kyo | L*er* | Sha |
| ***TIM22-2*** | An-1 | 69.0 | 37.9 | 26.7 | 29.9 | 55.0 | 21.3 | 64.7 | 18.6 |
| | C24 | 12.4 | 82.3 | 15.4 | 27.3 | 43.3 | 25.2 | 37.2 | 22.9 |
| | Col-0 | 18.7 | 25.6 | 55.7 | 57.8 | 27.6 | 42.7 | 20.7 | 18.2 |
| | Cvi-0 | 10.4 | 21.0 | 10.8 | 32.4 | 13.3 | 6.3 | 10.6 | 18.2 |
| | Eri-1 | 34.2 | 40.5 | 13.8 | 28.4 | 61.3 | 17.3 | 17.9 | 20.6 |
| | Kyo | 13.4 | 26.7 | 39.0 | 33.0 | 16.5 | 69.1 | 26.4 | 34.9 |
| | L*er* | 30.3 | 53.4 | 19.8 | 17.7 | 20.5 | 17.1 | 60.0 | 12.2 |
| | Sha | 22.7 | 27.7 | 20.7 | 55.4 | 17.5 | 54.1 | 28.5 | 54.3 |

**Supplementary Table 3.** The observed and expected frequencies of allele pair combinations of *TIM22-1* and *TIM22-2* in the AMPRIL subpopulation EGGE.

| | ***Observed*** | ***TIM22-1*** | | | |
|---|---|---|---|---|---|
| | | An-1 | Col-0 | Cvi-0 | L*er* |
| ***TIM22-2*** | An-1 | 26 | 21 | 20 | 11 |
| | Col-0 | 13 | 8 | 2 | 18 |
| | Cvi-0 | 1 | 0 | 12 | 1 |
| | L*er* | 4 | 8 | 12 | 19 |
| | ***Expected*** | ***TIM22-1*** | | | |
| | | An-1 | Col-0 | Cvi-0 | L*er* |
| ***TIM22-2*** | An-1 | 19.5 | 16.4 | 20.4 | 21.7 |
| | Col-0 | 10.3 | 8.6 | 10.7 | 11.4 |
| | Cvi-0 | 3.5 | 2.9 | 3.7 | 3.9 |
| | L*er* | 10.8 | 9.0 | 11.2 | 12.0 |

**Supplementary Table 4.** Loss-of-function (LoF) variation, expression (Exp.) and promoter methylation (Met.) of *TIM22-1* and *TIM22-2* in the AMPRIL parental genomes.

| | ***TIM22-1*** | | | | ***TIM22-2*** | | | |
|---|---|---|---|---|---|---|---|---|
| **Acc.** | **Gene ID** | **LoF** | **Exp.** | **Met.** | **Gene ID** | **LoF** | **Exp.** | **Met.** |
| An-1 | ATAN-1G29450 | - | + | - | ATAN-3G20030 | - | + | - |
| C24 | ATC24-1G29460 | - | + | - | ATC24-3G20040 | - | + | - |
| Col-0 | AT1G18320 | STG | + | - | AT3G10110 | - | + | - |
| Cvi-0 | ATCVI-1G29600 | - | + | - | ATCVI-3G19910 | - | - | + |
| Eri-1 | | STG | No RNA-seq data | - | ATERI-3G20040 | - | No RNA-seq data | - |
| Kyo | ATKYO-1G29480 | - | No RNA-seq data | - | ATKYO-3G20070 | - | No RNA-seq data | - |
| Ler | | STG | + | - | ATLER-3G19980 | - | + | - |
| Sha | | STG | + | - | ATSHA-3G25560 | - | + | - |

STG: stop-codon gain SNP

**Supplementary Table 5.** Loss-of-function (LoF) variation, expression (Exp.) and promoter methylation (Met.) of *HPA2* and *HPA1* in the AMPRIL parental genomes.

| | ***HPA2*** | | | | ***HPA1*** | | | |
|---|---|---|---|---|---|---|---|---|
| **Acc.** | **Gene ID** | **LoF** | **Exp.** | **Met.** | **Gene ID** | **LoF** | **Exp.** | **Met.** |
| An-1 | - | STG | + | - | ATAN-5G20070 | - | + | - |
| C24 | ATC24-1G85380 | - | - | + | ATC24-5G20050 | - | + | - |
| Col-0 | AT1G71920 | - | - | + | AT5G10330 | - | + | - |
| Cvi-0 | ATCVI-1G86350 | - | + | - | - | - | - | - |
| Eri-1 | - | STG | No RNA-seq data | - | ATERI-5G20170 | - | No RNA-seq data | - |
| Kyo | - | STG | No RNA-seq data | - | ATKYO-5G20130 | - | No RNA-seq data | - |
| L*er* | - | STG | + | - | ATLER-5G20130 | - | + | - |
| Sha | - | STG | + | - | ATSHA-5G20120 | - | + | - |

STG: stop-codon gain SNP

**Supplementary Table 6.** The observed and expected frequency of allele pair combinations of *HPA2* and *HPA1* in whole AMPRIL population

| ***Observed*** | | ***HPA2*** | | | | | | | |
|---|---|---|---|---|---|---|---|---|---|
| | | An-1 | C24 | Col-0 | Cvi-0 | Eri-1 | Kyo | Ler | Sha |
| ***HPA1*** | An-1 | 88 | 29 | 16 | 18 | 46 | 20 | 31 | 24 |
| | C24 | 26 | 68 | 31 | 24 | 32 | 25 | 46 | 21 |
| | Col-0 | 24 | 14 | 69 | 33 | 16 | 40 | 14 | 19 |
| | Cvi-0 | 2 | 1 | 14 | 80 | 2 | 11 | 2 | 14 |
| | Eri-1 | 59 | 44 | 30 | 19 | 78 | 16 | 21 | 24 |
| | Kyo | 15 | 24 | 65 | 21 | 25 | 83 | 10 | 40 |
| | Ler | 52 | 59 | 33 | 23 | 18 | 29 | 47 | 24 |
| | Sha | 24 | 19 | 19 | 34 | 17 | 40 | 13 | 59 |

| ***Expected*** | | ***HPA2*** | | | | | | | |
|---|---|---|---|---|---|---|---|---|---|
| | | An-1 | C24 | Col-0 | Cvi-0 | Eri-1 | Kyo | Ler | Sha |
| ***HPA1*** | An-1 | 79 | 28 | 20 | 30 | 38 | 19 | 35 | 22 |
| | C24 | 24 | 68 | 35 | 32 | 42 | 19 | 36 | 19 |
| | Col-0 | 22 | 15 | 66 | 37 | 17 | 40 | 14 | 18 |
| | Cvi-0 | 12 | 8 | 24 | 33 | 9 | 14 | 8 | 19 |
| | Eri-1 | 54 | 44 | 32 | 31 | 67 | 21 | 16 | 27 |
| | Kyo | 19 | 27 | 56 | 21 | 25 | 79 | 13 | 42 |
| | Ler | 56 | 50 | 23 | 34 | 19 | 28 | 50 | 26 |
| | Sha | 25 | 18 | 21 | 35 | 17 | 45 | 12 | 52 |

**Supplementary Table 7.** The observed and expected frequency of allele pair combinations of *HPA2* and *HPA1* in subpopulation ABBA population

| ***Observed*** | | ***HPA2*** | | | |
|---|---|---|---|---|---|
| | | Col-0 | Cvi-0 | Kyo | Sha |
| ***HPA1*** | Col-0 | 9 | 7 | 6 | 3 |
| | Cvi-0 | 9 | 8 | 8 | 5 |
| | Kyo | 16 | 10 | 9 | 6 |
| | Sha | 5 | 9 | 7 | 9 |

| ***Expected*** | | ***HPA2*** | | | |
|---|---|---|---|---|---|
| | | Col-0 | Cvi-0 | Kyo | Sha |
| ***HPA1*** | Col-0 | 7.2 | 6.3 | 7.4 | 4.2 |
| | Cvi-0 | 8.6 | 7.5 | 8.8 | 5.1 |
| | Kyo | 14.6 | 12.8 | 15.0 | 8.6 |
| | Sha | 8.6 | 7.5 | 8.8 | 5.1 |

**Supplementary Table 8.** The observed and expected frequency of allele pair combinations of *HPA2* and *HPA1* in subpopulation EFFE population

| ***Observed*** | | ***HPA2*** | | | |
|---|---|---|---|---|---|
| | | Col-0 | Cvi-0 | Kyo | Sha |
| ***HPA1*** | Col-0 | 17 | 4 | 18 | 16 |
| | Cvi-0 | 3 | 5 | 3 | 4 |
| | Kyo | 18 | 11 | 17 | 15 |
| | Sha | 14 | 3 | 16 | 8 |

| ***Expected*** | | ***HPA2*** | | | |
|---|---|---|---|---|---|
| | | Col-0 | Cvi-0 | Kyo | Sha |
| ***HPA1*** | Col-0 | 16.6 | 7.4 | 17.3 | 13.8 |
| | Cvi-0 | 4.5 | 2.0 | 4.7 | 3.8 |
| | Kyo | 18.4 | 8.2 | 19.2 | 15.3 |
| | Sha | 12.4 | 5.5 | 12.9 | 10.3 |

**Supplementary Table 9.** Pseudo-heterozygous variants identified in *HPA2* and *HPA1* by RNA-seq and methylation data.

| **Gene** | **Variation ID** | **Chr.** | **Position** | **Reference allele** | **Alternative allele** | **Functional effect** | **Number of accessions** | **Variation could be assigned to *HPA2* or *HPA1*** |
|---|---|---|---|---|---|---|---|---|
| *HPA2* | LoF-100 | chr1 | 27,067,175 | T | TCAGGGCTCTC | FS | 1 | *HPA2* |
| *HPA2* | LoF-110 | chr1 | 27,067,231 | C | T | STG | 2 | *HPA2* |
| Both | LoF-120 | chr1 | 27,067,387 | ATTTT | ATTT | FS | 3 | *HPA2* |
| Both | LoF-120 | chr5 | 3,249,634 | TAAAA | TAAA | FS | 2 | *HPA2* |
| Both | LoF-130 | chr1 | 27,067,518 | C | T | STG | 2 | Undet. |
| Both | LoF-130 | chr5 | 3,249,508 | G | A | STG | 2 | Undet. |
| *HPA1* | LoF-140 | chr5 | 3,249,497 | TC | TCC | FS | 1 | Undet. |
| Both | LoF-150 | chr1 | 27,067,590 | G | A | spDon | 1 | Undet. |
| Both | LoF-150 | chr5 | 3,249,436 | C | T | spDon | 1 | Undet. |
| *HPA1* | LoF-160 | chr1 | 27,067,591 | C | A | spDon | 1 | Undet. |
| Both | LoF-170 | chr1 | 27,067,785 | CT | C | FS | 210 | *HPA2* |
| Both | LoF-170 | chr5 | 3,249,239 | GA | G | FS | 212 | *HPA2* |
| Both | LoF-180 | chr1 | 27,067,787 | CAA | CA | STG | 1 | Undet. |
| Both | LoF-180 | chr5 | 3,249,236 | CTT | CT | FS | 1 | Undet. |
| Both | LoF-190 | chr1 | 27,067,813 | CATACTTGTA | CA | FS | 23 | Undet. |
| Both | LoF-190 | chr5 | 3,249,203 | CTACAAGTAT | CT | FS | 16 | Undet. |
| *HPA1* | LoF-200 | chr5 | 3,249,064 | C | A | STG | 1 | Undet. |
| *HPA1* | LoF-210 | chr5 | 3,249,052 | CT | C | FS | 1 | *HPA2* |
| Both | LoF-220 | chr1 | 27,067,987 | CTTT | CTT | FS | 1 | *HPA2* |
| Both | LoF-220 | chr5 | 3,249,035 | GAAA | GAA | FS | 1 | *HPA2* |
| Both | LoF-230 | chr1 | 27,068,021 | TGG | TG | FS | 12 | *HPA2* |
| Both | LoF-230 | chr5 | 3,249,002 | TCC | TC | FS | 13 | *HPA2* |
| Both | LoF-240 | chr1 | 27,068,037 | A | T | STG | 1 | Undet. |
| Both | LoF-240 | chr5 | 3,248,989 | T | A | STG | 1 | Undet. |
| Both | LoF-250 | chr1 | 27,068,199 | G | T | STG | 1 | *HPA2* |
| Both | LoF-250 | chr5 | 3,248,827 | C | A | STG | 1 | *HPA2* |
| Both | LoF-260 | chr1 | 27,068,344 | G | A | spAcc | 1 | *HPA2* |
| Both | LoF-260 | chr5 | 3,248,682 | C | T | spAcc | 1 | *HPA2* |
| Both | LoF-270 | chr1 | 27,068,433 | G | T | STG | 1 | Undet. |
| Both | LoF-270 | chr5 | 3,248,593 | C | A | STG | 1 | Undet. |
| Both | LoF-280 | chr1 | 27,068,634 | GAA | GAAA | FS | 1 | *HPA2* |
| Both | LoF-280 | chr5 | 3,248,392 | C | A | STG | 1 | *HPA2* |
| Both | LoF-290 | chr1 | 27,068,644 | G | A | STG | 30 | Undet. |
| Both | LoF-290 | chr5 | 3,248,382 | C | T | STG | 16 | Undet. |
| Both | LoF-300 | chr1 | 27,068,655 | C | T | STG | 585 | *HPA2* |
| Both | LoF-300 | chr5 | 3,248,371 | G | A | STG | 405 | *HPA2* |
| *HPA1* | LoF-310 | chr5 | 3,248,194 | C | A | STG | 1 | *HPA2* |
| Both | LoF-320 | chr1 | 27,068,893 | TGG | TG | FS | 3 | Undet. |
| Both | LoF-320 | chr5 | 3,248,130 | ACC | AC | FS | 4 | Undet. |

Both: variation was found in both *HPA2* and *HPA1*; FS: frameshift; STG: premature stop-codon gain; spAcc: splice acceptor site damage; spDon: splice donor site damage; Undet.: undetermined.

**Supplementary Table 10.** Number of accessions with different combinations of functional (FUN) or non-functional (nFun) *HPA* copies across the 1135 *A. thaliana* accessions of the 1001 Genomes Project. ("Pre" and "Abs" refer to the presence and absence of *HPA3/4* which could only be analyzed for their presence.)

| **Allele Combination** | ***HPA2*** | ***HPA1*** | ***HPA3*** | ***HPA4*** | **Number** |
|---|---|---|---|---|---|
| Allele Combination 1 | FUN | Abs | Abs | Abs | 3 |
| Allele Combination 2 | FUN | nFUN | Abs | Abs | 1 |
| Allele Combination 3 | nFUN | FUN | Abs | Abs | 661 |
| Allele Combination 4 | FUN | FUN | Abs | Abs | 266 |
| Allele Combination 5 | FUN | FUN | Pre | Abs | 48 |
| Allele Combination 6 | nFUN | FUN | Pre | Abs | 100 |
| Allele Combination 7 | FUN | nFUN | Pre | Abs | 1 |
| Allele Combination 8 | nFUN | nFUN | Pre | Abs | 13 |
| Allele Combination 9 | nFUN | FUN | Abs | Pre | 6 |
| Allele Combination 10 | nFUN | nFUN | Abs | Pre | 36 |

**Supplementary Table 11.** Number of accessions with different combinations of functional (FUN) or non-functional (nFUN) *TIM22* copies across 1135 *A, thaliana* accessions of the 1001 Genomes Project.

| | ***TIM22-1*** | ***TIM22-2*** | **Number** |
|---|---|---|---|
| Allele Combination 1 | FUN | FUN | 890 |
| Allele Combination 2 | FUN | nFUN | 4 |
| Allele Combination 3 | nFUN | FUN | 225 |
| Allele Combination 4 | nFUN | nFUN | 16 |

**Supplementary Table 12.** Loss-of-function (LoF) variation, expression (Exp.) and promoter methylation (Met.) of *FOLT1* and *FOLT2* in the AMPRIL parental genomes.

| | ***FOLT1*** | | | | ***FOLT2*** | | | |
|---|---|---|---|---|---|---|---|---|
| **Acc.** | **Gene ID** | **LoF** | **Exp.** | **Met.** | **Gene ID** | **LoF** | **Exp.** | **Met.** |
| An-1 | ATAN-5G89880 | - | + | - | ATAN-4G54430 | - | + | - |
| C24 | ATC24-5G86520 | - | - | + | ATC24-4G57120 | - | + | - |
| Col-0 | AT5G66380 | - | + | - | - | - | - | - |
| Cvi-0 | ATCVI-5G87930 | - | + | - | ATCVI-4G51750 | - | + | - |
| Eri-1 | ATERI-5G89000 | - | No RNA-seq data | - | ATERI-4G53100 | - | No RNA-seq data | - |
| Kyo | ATKYO-5G89250 | - | No RNA-seq data | - | in a contig | - | No RNA-seq data | - |
| L*er* | ATLER-5G86770 | - | + | - | ATLER-4G54710 | - | + | - |
| Sha | ATSHA-5G91420 | - | - | + | ATSHA-4G55130 | - | + | - |

**Supplementary Table 13.** The number of accessions with different combinations of functional (FUN), non-functional (nFUN) and truncated (trun.) alleles of *FOLT* across 1,135 *A. thaliana* accessions from the 1001 Genomes Project.

| **Allele Combination** | ***FOLT1*** | ***FOLT2*** | ***FOLT2* trun.** | **Number** |
|---|---|---|---|---|
| Allele Combination 1 | FUN | Abs | Abs | 195 |
| Allele Combination 2 | FUN | FUN | Abs | 865 |
| Allele Combination 3 | nFUN | FUN | Abs | 29 |
| Allele Combination 4 | nFUN | FUN | trExon1-2 | 14 |
| Allele Combination 5 | nFUN | FUN | trExon1-6 | 1 |
| Allele Combination 6 | nFUN | FUN | trExon1-2 + trExon1-6 | 31 |

Note: trExon1-2 and trExon1-6 indicate that the truncated copy contains the first two or first six exons of *FOLT2*, respectively. “Abs” refers to the absence of intact or truncated *FOLT2*.

**Supplementary Table 14.** The number of accessions with different combinations of functional alleles of *TAD3* across 1,135 *A. thaliana* accessions from the 1001 Genomes Project

| Functional Allele | Number |
|---|---|
| *TAD3-1* | 971 |
| *TAD3-2* | 150 |
| *TAD3-1, TAD3-2* | 14 |

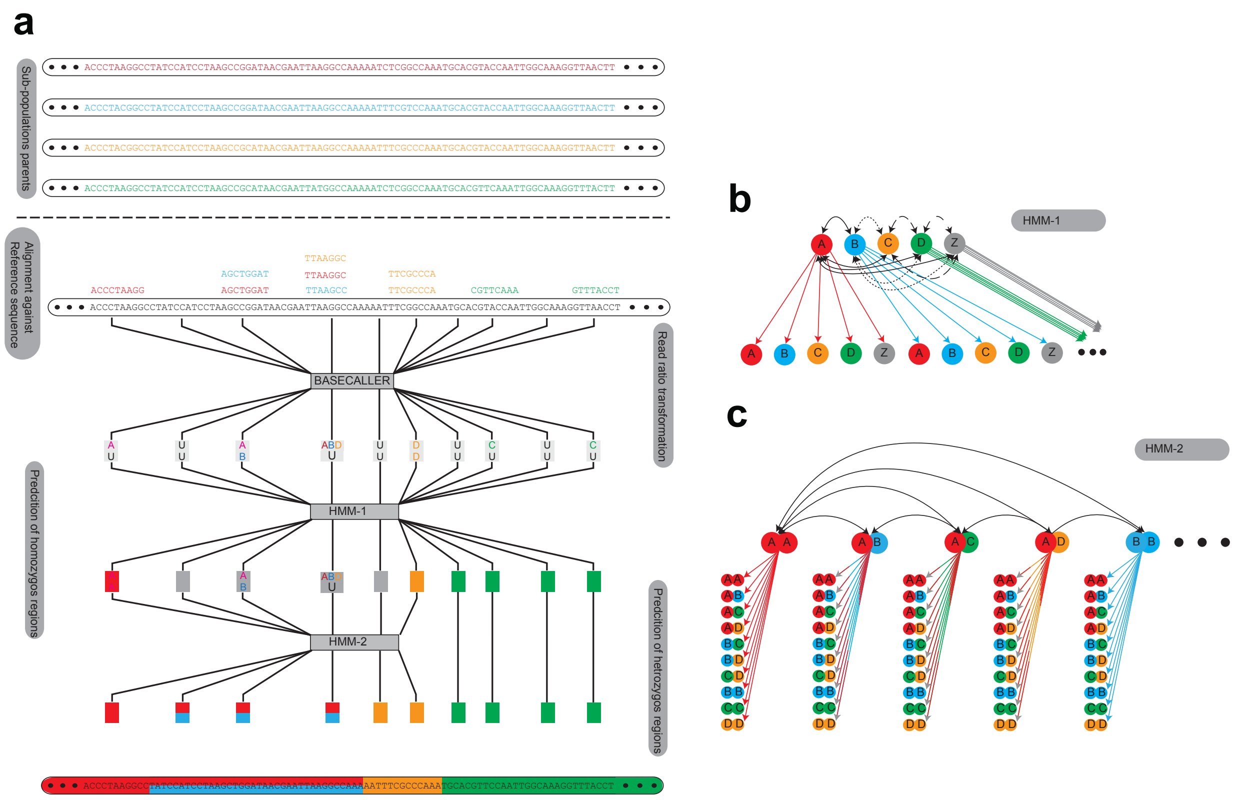


**Supplementary Fig. 1 Workflow and HMMs genotyping AMPRIL population**. **(a)** Schematic workflow of genotyping. The allele from short read sequencing were translated into genotype calls considering the diploid nature of the genomes. **(b, c)** Hidden State Models of the two HMMs, which were used for genotyping of the AMPRILs. The HMM-Model-1 was used to assign homozygous genotypes, while it did not resolve heterozygous regions (collectively assigned as Z regions). The more complex HMM-Model-2 was then used locally to resolve Z regions into heterozygous states.

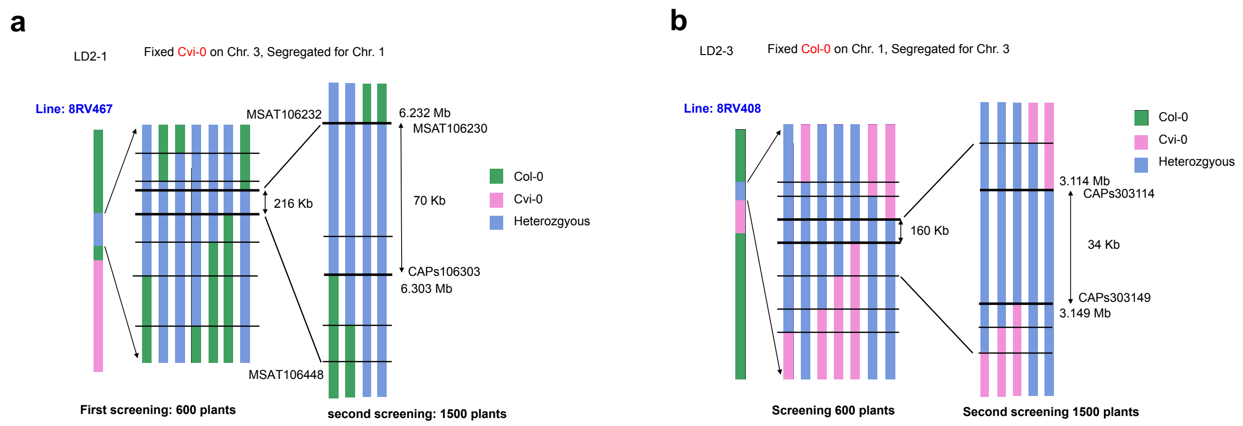


**Supplementary Fig. 2** Two rounds of fine-mapping using residual heterozygosity among heterogeneous inbred families from a Cvi-0 x Col-0 RIL population screened genotypically at **(a)** LD2.1 or **(b)** LD2.3 loci.

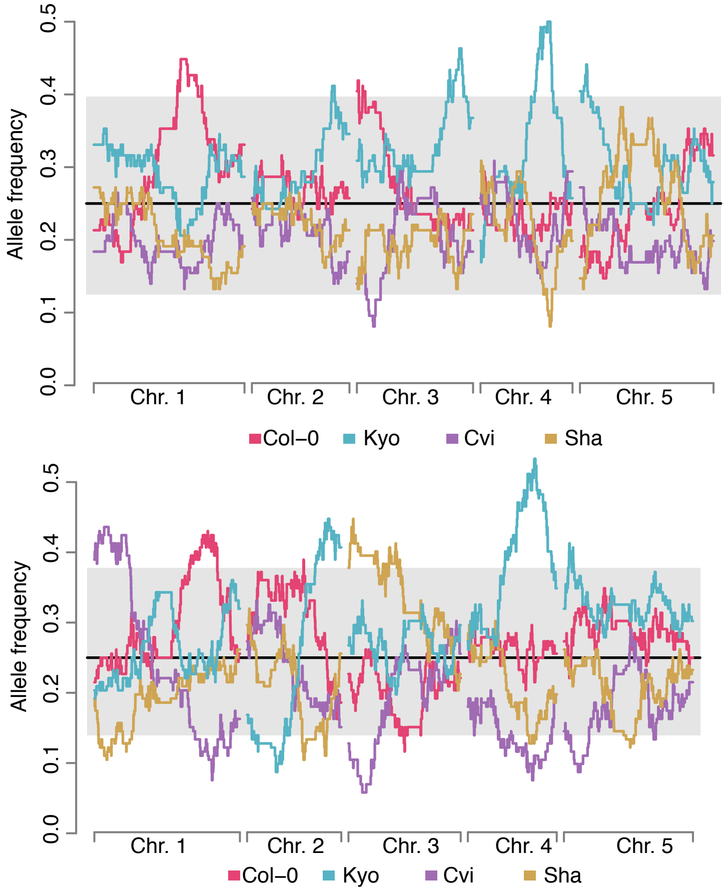


**Supplementary Fig. 3** Allele frequency distribution along the chromosome in subpopulation ABBA (above) and EFFE (below)

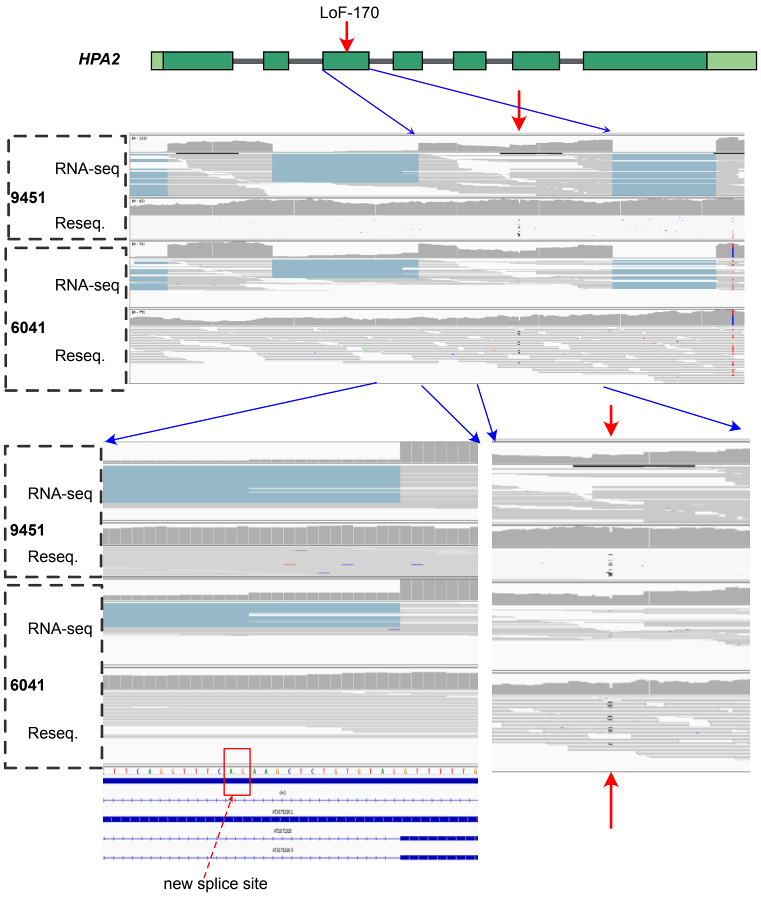


**Supplementary Fig. 4 Snapshot of IGV[4] of *A. thaliana* accessions with the frameshift mutation (LoF-170) in *HPA2* which was rescued by alternative splicing**. The position of heterozygous variant of LoF-170 is marked with red arrow. In both DNA short read resequencing alignments (Reseq.) of accession Spro 2 (accession ID: 9451) and Lis-3 (accession ID: 6041) the LoF-170 is found. The mutation is however only found in parts of the reads as reads from *HPA1* and *HPA2* are aligned repetitively. However, in RNA-seq read mapping no LoF-170 is found in Spro 2. This suggests that LoF-170 occurred in *HPA2,* because *HPA2* is most likely not expressed due to hypermethylation of its promoter. In contrast, the LoF-170 is still found in RNA-seq reads of accession Lis-3. While this could imply that *HPA2* is expressed, the mutation is linked to an alternative splicing site (indicated by the red dashed arrow) which can complement this frameshift of LoF-170. (Blue regions in RNA-seq read mapping indicate the introns.)

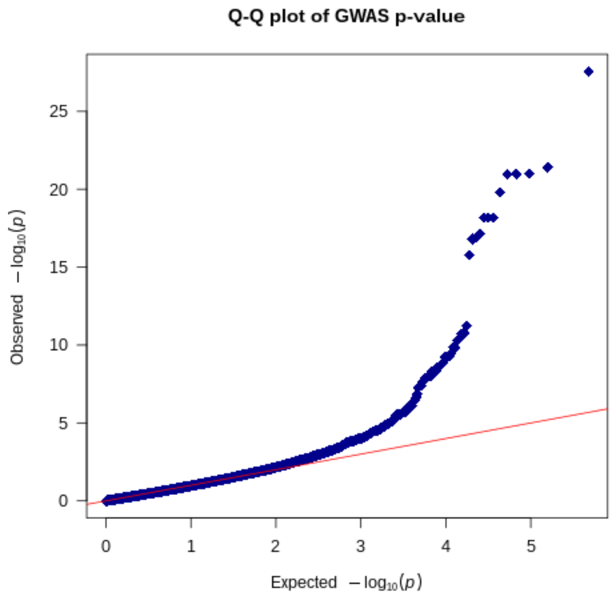


**Supplementary Fig. 5. QQ-plot of the GWAS using the absence/presence of functional *HPA* copies as phenotype**.

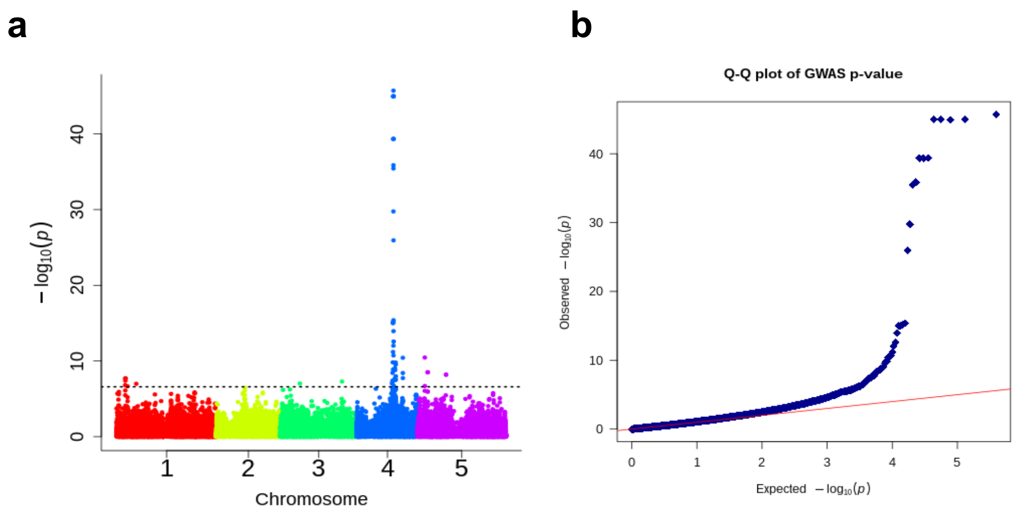


**Supplementary Fig. 6. Manhattan plot (a) and QQ-plot (b) of the GWAS using the absence/presence of functional *HPA* copies as phenotype**. In contrast to the same approach as shown in Figure 4, samples with *HPA3* were excluded from this analysis.

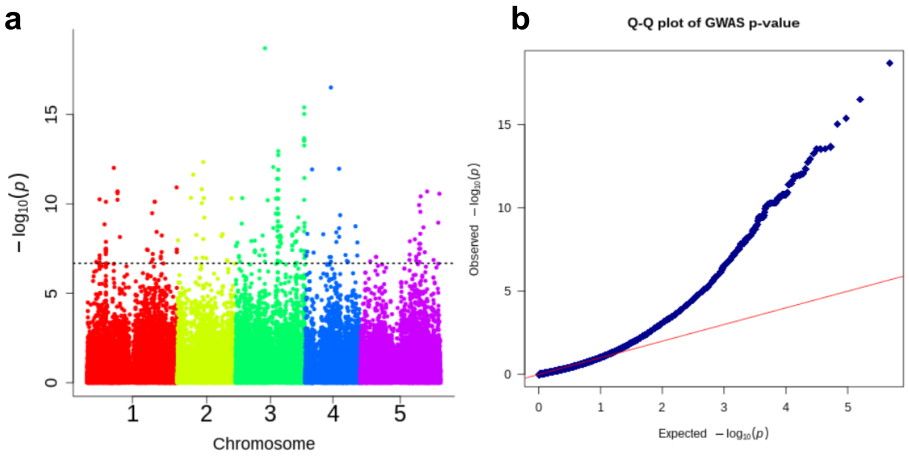


**Supplementary Fig. 7 Manhattan plot (a) and QQ-plot (b) of the GWAS using absence/presence of functional *TIM22* copies as phenotype.**

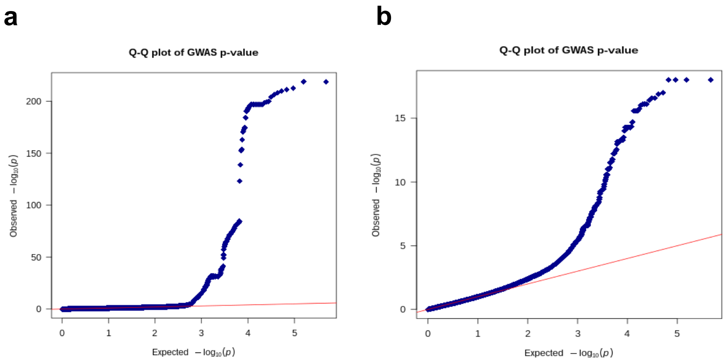


**Supplementary Fig. 8 QQ-plot of GWAS for the phenotype of presence or absence of functional *TAD3-1* copy (a) and *FOLT-1* copy (b).**

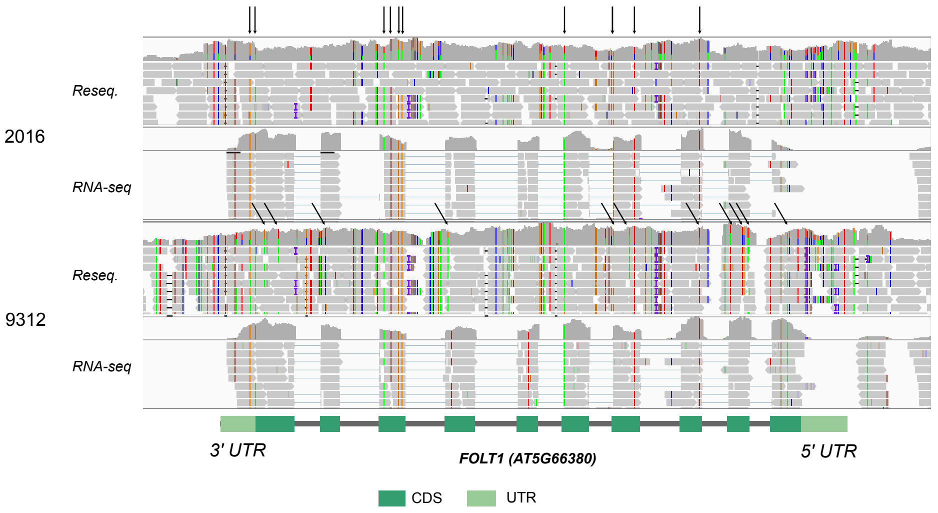


**Supplementary Fig. 9 Snapshot of IGV for examples of accessions with expression-silenced *FOLT1* and expression-active *FOLT2*.** The heterozygous variants in resequencing reads are marked with arrows, which appear homozygous in the RNA-seq data. Two accessions MNF-Pin-39 (accession ID: 2016) and Ullapool-8 (accession ID: 9312) are used as the examples.

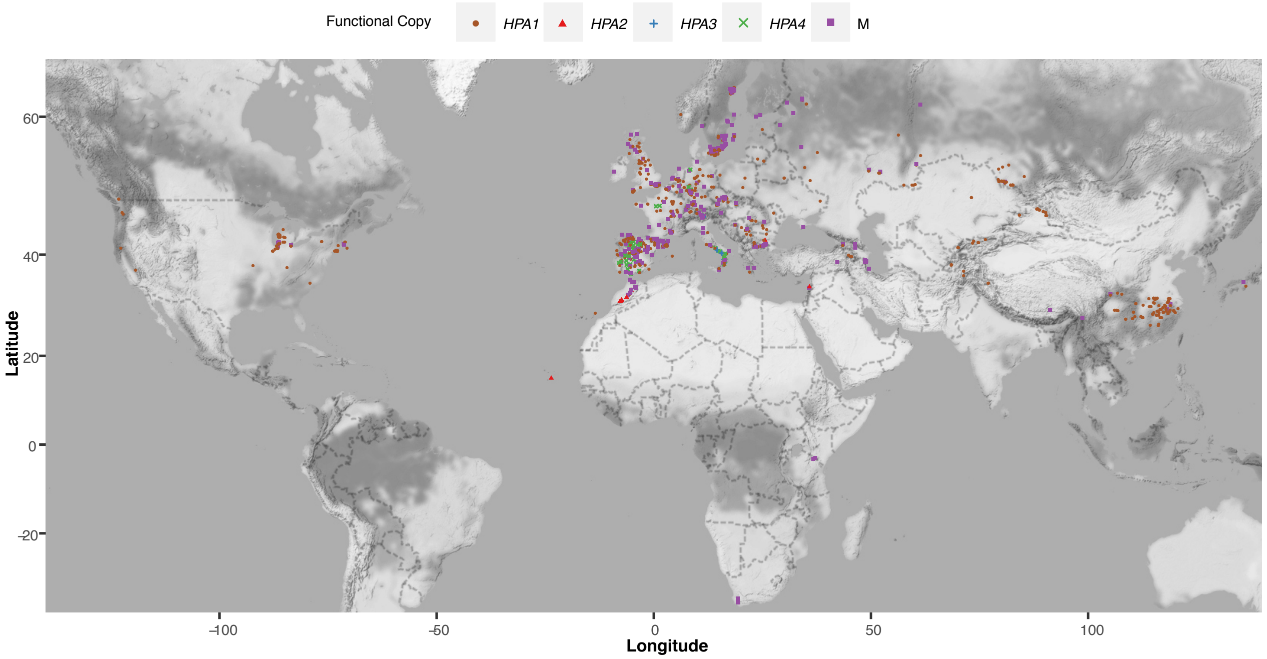


**Supplementary Fig. 10 Geographic distribution of functional copies of *HPA* across 1135 *A. thaliana* accessions from 1001 Genomes Project, 75 African accessions and 118 Chinese accessions**. M: multiple functional copies, Other accessions only with single functional copy are shown.

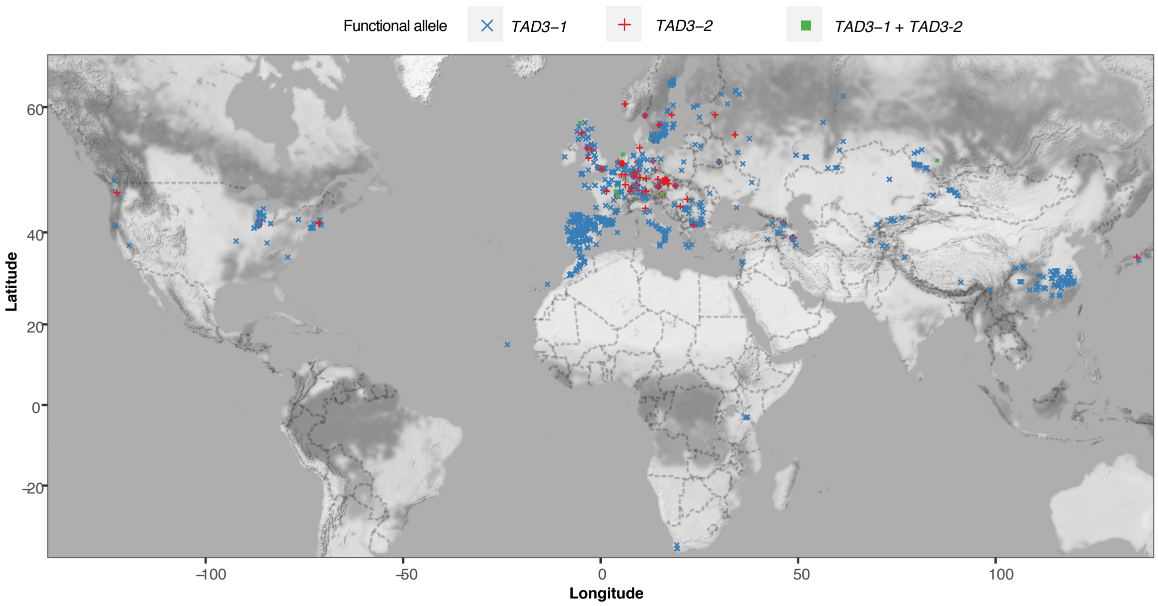


**Supplementary Fig. 11 Geographic distribution of functional copies of *TAD3* across 1135 *A. thaliana* accessions from 1001 Genomes Project, 75 African accessions and 118 Chinese accessions.**

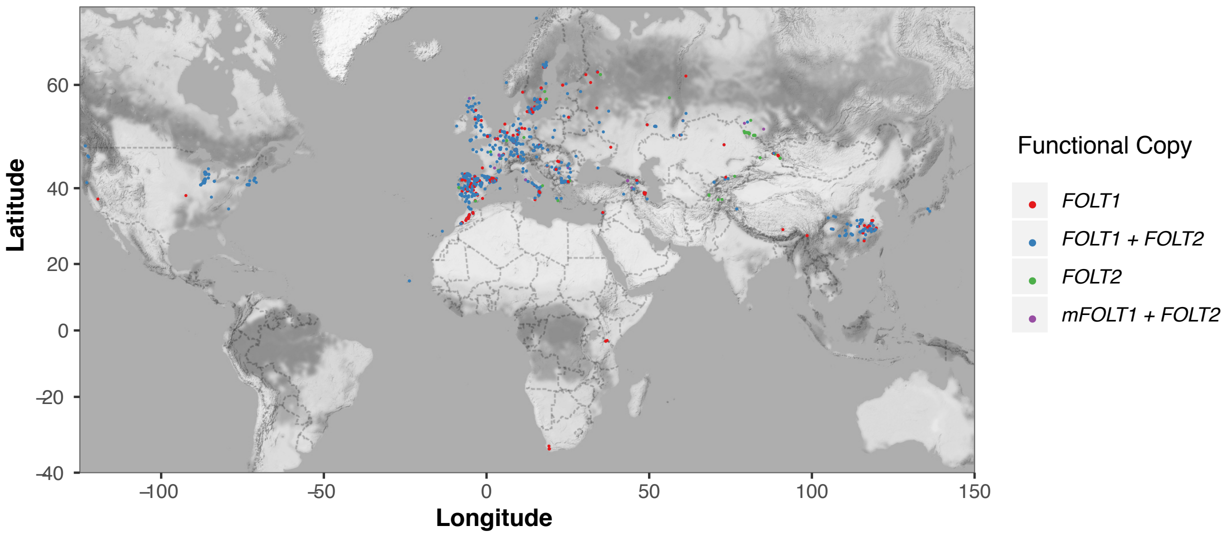


**Supplementary Fig. 12 Geographic distribution of functional copies of *FOLT* across 1135 *A. thaliana* accessions from 1001 Genomes Project, 75 African accessions and 118 Chinese accessions**. *mFOLT1* + *FOLT2*: accessions with a functional copy *FOLT2* and an unexpressed copy *FOLT1* but without truncated copy of *FOLT*.

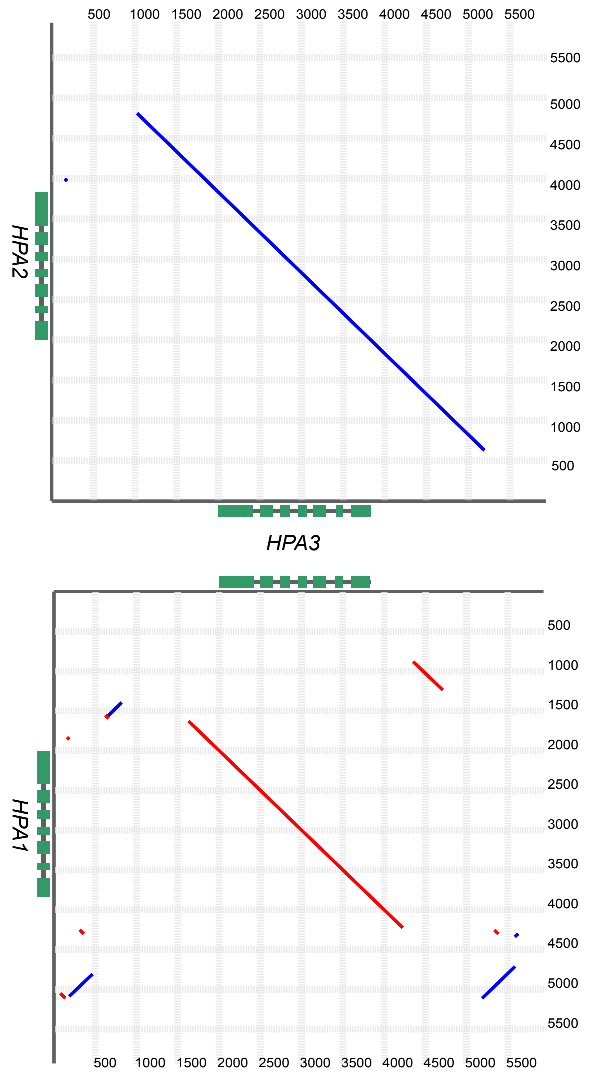


**Supplementary Fig. 13 Sequence alignment between *HPA3* and two other *HPA* copies, *HPA1* and *HPA2.***

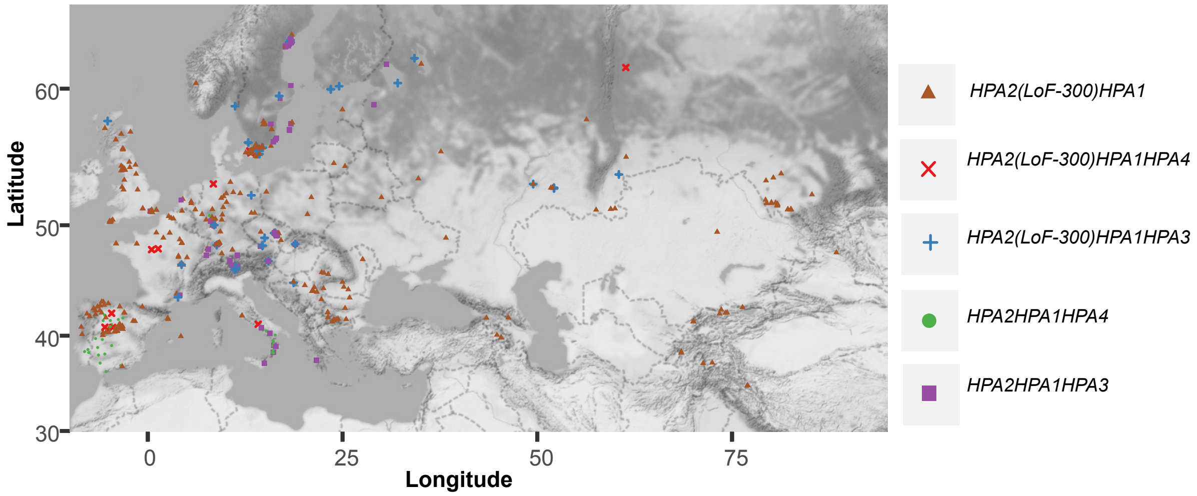


**Supplementary Fig. 14 Geographic distribution of Eurasian *A. thaliana* accessions with loss-of-function (LoF) variants in *HPA2* across accessions with the copy of *HPA3* or *HPA4.***

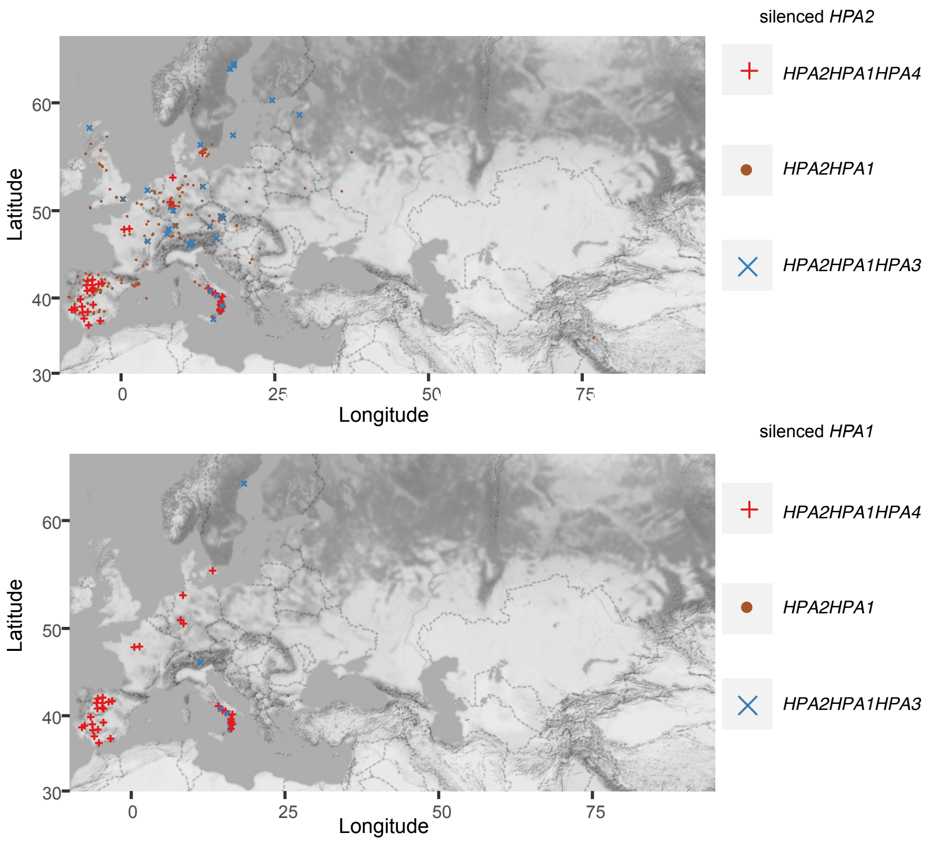


**Supplementary Fig. 15 Geographic distribution of *A. thaliana* accessions with different *HPA* copies and methylated promoter in *HPA2* (upper panel) or *HPA1* (lower panel).**

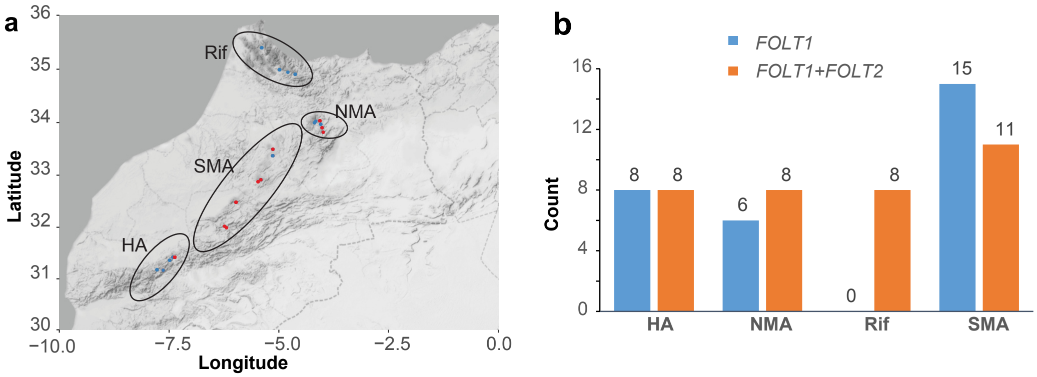


**Supplementary Fig. 16 Distribution (a) and number (b) of *FOLT* copies across African accessions.**

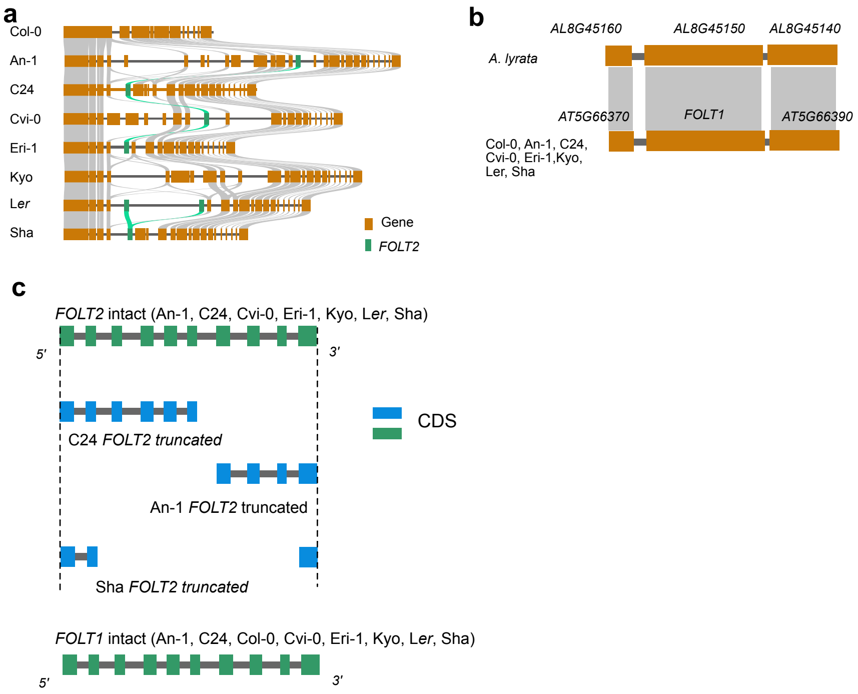


**Supplementary Fig. 17 Gene structure of *FOLT* copies in the eight AMPRIL parental genomes as well as in *A. lyrata*. (a)** Gene arrangements around the *FOLT2* locus in the AMPRIL eight parental genomes. Gray links indicate orthologous genes. The *FOLT2* gene in Kyo is in a short unanchored contig. **(b)** The *FOLT1* gene in *A. lyrata* and eight *A. thaliana* ecotypes. All of them share syntenic gene arrangement (as indicated by the gray bars). **(c)** The *FOLT2* and *FOLT1* gene structure as observed in the AMPRIL.
